# Defective ribosome assembly impairs leukemia stem cell function in a murine model of acute myeloid leukemia

**DOI:** 10.1101/2023.01.22.525120

**Authors:** Daniel Sjövall, Sudip Ghosh, Jenny Hansson, Carolina Guibentif, Pekka Jaako

**Author notes:** **Corresponding author:** Pekka Jaako, Sahlgrenska Center for Cancer Research, University of Gothenburg, Box 425, SE-40530, Gothenburg, Sweden.

## Abstract

Despite the advanced understanding of disease mechanisms, the current therapeutic regimen fails to cure most patients with acute myeloid leukemia (AML). In the present study, we address the role of protein synthesis control in AML leukemia stem cell (LSC) function and leukemia propagation. We apply a murine model of mixed-lineage leukemia-rearranged AML to demonstrate that LSCs are characterized by high global protein synthesis rate. Using a genetic model that permits inducible and graded regulation of ribosomal subunit joining, we show that defective ribosome assembly leads to a significant survival advantage by selectively eradicating LSCs but not normal hematopoietic stem and progenitor cells. Finally, transcriptomic and proteomic analyses identify a rare subset of LSCs with immature stem cell signature and high ribosome content that promotes the resistance to defective ribosome assembly. Collectively, our study unveils a critical requirement of high protein synthesis rate for LSC function and highlights ribosome assembly as a therapeutic target in AML.

## INTRODUCTION

Acute myeloid leukemia (AML) is a heterogenous hematologic malignancy characterized by clonal expansion of aberrant myeloid precursor cells and bone marrow (BM) failure. Despite significant advances in understanding the AML disease mechanisms, the standard therapeutic regimen has remained practically unchanged for more than four decades^1^. This regimen induces high rates of remission, but in most cases fails to eradicate leukemia stem cells (LSCs) that ultimately underlie disease relapse. There is therefore an unmet clinical need for novel therapeutic strategies that selectively target LSCs.

Most patients with AML harbor mutations in genes that encode signal transduction pathway components^2^. These mutations typically arise late during disease progression and promote the conversion of pre-leukemic myeloid progenitor cells into AML LSCs^3,4^. Most notably, approximately one third of all AML cases show mutations in the gene encoding FLT3 receptor tyrosine kinase that result in constitutive activation of PI3K/AKT/mTOR and RAS/MAPK pathways^2^. However, despite the development of effective inhibitors, therapeutic targeting of FLT3 has remained challenging due to the emergence of subclones with RAS pathway mutations or on-target *FLT3* mutations^5^. Moreover, phase I/II trials targeting mTOR signaling have generally failed to show meaningful responses in AML^6^.

The mutations that activate oncogenic signal transduction pathways in AML are largely mutually exclusive, suggesting that they drive leukemia by converging on common downstream mechanisms^2^. An accumulating body of evidence suggests that most oncogenic signaling pathways regulate ribosome biogenesis and protein synthesis^7,8^. These findings suggest a critical role for protein synthesis control in LSC biology that may represent a broad therapeutic vulnerability in AML. Consistently, the protein synthesis inhibitor homoharringtonine has been used in the treatment of hematologic malignancies in China, and it was recently approved by the United States Food and Drug Administration for the treatment of tyrosine kinase inhibitor-resistant chronic myeloid leukemia^9^. Moreover, the RNA polymerase I inhibitor CX-5461, which inhibits ribosomal RNA transcription, outperforms standard chemotherapies in mouse models of AML and it is currently being evaluated in several phase I clinical trials^10^.

Despite the promise of protein synthesis machinery as a target in AML, the cellular and molecular mechanisms that define the response of leukemia cells to ribosomal insults remain incompletely understood. In particular, the difficulty to model ribosomal dysfunction in mice has presented a significant barrier for the validation of the biological and therapeutic role of protein synthesis machinery in AML in vivo. In the present study, we apply a recently generated transgenic mouse model of defective ribosome assembly to demonstrate a critical and selective requirement of high protein synthesis rate for LSC function and leukemia propagation, highlighting the protein synthesis machinery as a therapeutic target in AML.

## MATERIAL AND METHODS

### Animal studies

All animal procedures were approved by the Ethical Committee for Animal Experiments at the University of Gothenburg. Detailed information on mouse strains and experimental procedures can be found in the Supplemental Information.

### Transplantation assays

MLL-AF9 AML model systems were generated by transplanting one million bulk-transduced cells (CD45.2) together with 0.5 million CD45.1 BM cells into lethally irradiated (2x 475 rad, X-ray source) CD45.1 recipient mice. All experiments were performed by transplanting 50,000 or 100,000 BM cells from serial transplanted, terminally sick mice into sublethally irradiated (475 rad) recipient mice. The C57BL/6J and [M2-rtTA/M2-rtTA][EIF6/+] AML were of male origin. The [M2-rtTA/M2-rtTA][EIF6/EIF6] AML was of female origin. Both female and male mice were used as hosts for leukemia grafts. Non-competitive transplantation experiment was performed by injecting three million unfractionated [M2-rtTA/M2-rtTA][EIF6/EIF6] BM cells from a male donor (CD45.2) into lethally irradiated (2x 475 rad) female recipient mice (CD45.1).

### Bone marrow harvest

BM was harvested by crushing the hips, femurs and tibias in FACS buffer (PBS supplemented with 2% fetal calf serum (FCS; Merck) and 2 mM EDTA (Merck). Isolated cells were passed through a 70 μm cell strainer (Thermo Fisher Scientific).

### Flow cytometry

Five million BM cells were stained with antibodies in FACS buffer for 30 min on ice and protected from light. Antibodies used are listed in **Supplementary Table 1**. Propidium iodide (Invitrogen) was used to label dead cells. Flow cytometry was performed using BD LSRFortessa, BD LSR II and BD FACSAria Fusion flow cytometers (BD Biosciences). Data analysis was performed using the FlowJo software (v10.8.1; BD Biosciences).

### Protein synthesis rate measurement

Mice were injected intraperitoneally with 200 μL of OP-Puro solution (3.4 mM; MedChemExpress, Jena Bioscience), and sacrificed one hour later. Three million BM cells were stained with antibodies against cell surface markers, and then fixed with 2% formaldehyde (Alfa Aesar) for 15 min on ice. Stained cells were permeabilized using PBS supplemented with 0.1% saponin (Merck) and 3% bovine serum albumin (Merck). The azide-alkyne reaction was performed using the Click-iT™ Cell Reaction Buffer Kit (Thermo Fisher Scientific) with Alexa Fluor™ 647 Azide (Thermo Fisher Scientific) at a final concentration of 2.5 μM.

### RNA sequencing and data analysis

BM cells were harvested from leukemia-engrafted mice after five days of Dox administration and 500,000 GFP+/c-Kit+ leukemia cells were collected using FACS. RNA extraction was performed using the RNeasy Mini Kit (Qiagen) and RNA-Integrity was assessed on the Bioanalyzer 2100 (Agilent; RIN ≥ 8 for all samples except Control-1 (7.33) and eIF6-4 (6.99). RNA libraries were constructed using TruSeq Stranded mRNA kit (Illumina). Pooled libraries were sequenced on NovaSeq 6000 system (Illumina). Data pre-processing was performed using the rnaseq pipeline^11^. *EIF6* and *MLL-AF9-GFP* transgene sequences were concatenated to *Mus musculus* GRCm38 reference genome. Differential expression analysis was performed using DESeq2 (v.1.34.0)^12^. Visualization of cell type enrichment of differentially expressed genes was performed using CellRadar (karlssonG.github.io/cellradar). Gene Set Enrichment Analysis (GSEA) was performed using v.4.1.0 software with default settings for small number of samples (“gene_set” permutation type)^13^.

### Statistical analyses

Student’s *t*-test was used to determine statistical significance, and two-tailed *P* values are shown. Data are presented as mean +/- standard deviation.

## RESULTS

### Leukemia stem cells show high global protein synthesis rate

Chromosomal rearrangements involving the *mixed-lineage leukemia 1* (*MLL1*; also known as *KMT2A*) gene generate powerful oncogenic fusion proteins that cause an aggressive and therapy-resistant subtype of AML^15^. To study protein synthesis control in MLL-rearranged AML, we transduced lineage-depleted BM cells with MLL-AF9-GFP retrovirus and transplanted them into congenic recipient mice to generate an aggressive AML **(Figure 1A)**. Previous studies have shown that leukemia-propagating LSCs in this model system account for 25-30% of leukemia cells in terminally sick mice and can be enriched to a high purity based on the presence of c-Kit cell surface receptor^16^. We could verify that c-Kit+ leukemia cells produced more colonies and showed higher proliferative potential compared with c-Kit- fraction **(Supplementary Figure 1)**. To quantify global protein synthesis rate in vivo, we performed O-propargyl-puromycin (OP-Puro) labeling experiments in leukemia-engrafted mice. OP-Puro is a puromycin analog that is incorporated into newly synthesized polypeptides, which can be quantified using flow cytometry **(Figure 1B)**^17^. These experiments showed that c-Kit+ LSCs represent one of the most highly-translating cell types in the BM, incorporating approximately six and three times more OP-Puro compared with c-Kit- leukemia cells and normal host-derived CD11b+ myeloid BM cells, respectively **(Figure 1B and 1C)**. In contrast to c-Kit+ LSCs, which showed a uniformly high OP-Puro incorporation rate, c-Kit- leukemia cells comprised a major OP-Puro^low^ subset and a minor OP-Puro^high^ subset. Taken together, these results demonstrate that c-Kit+ LSCs are characterized by high global protein synthesis rate.

**Figure 1.**
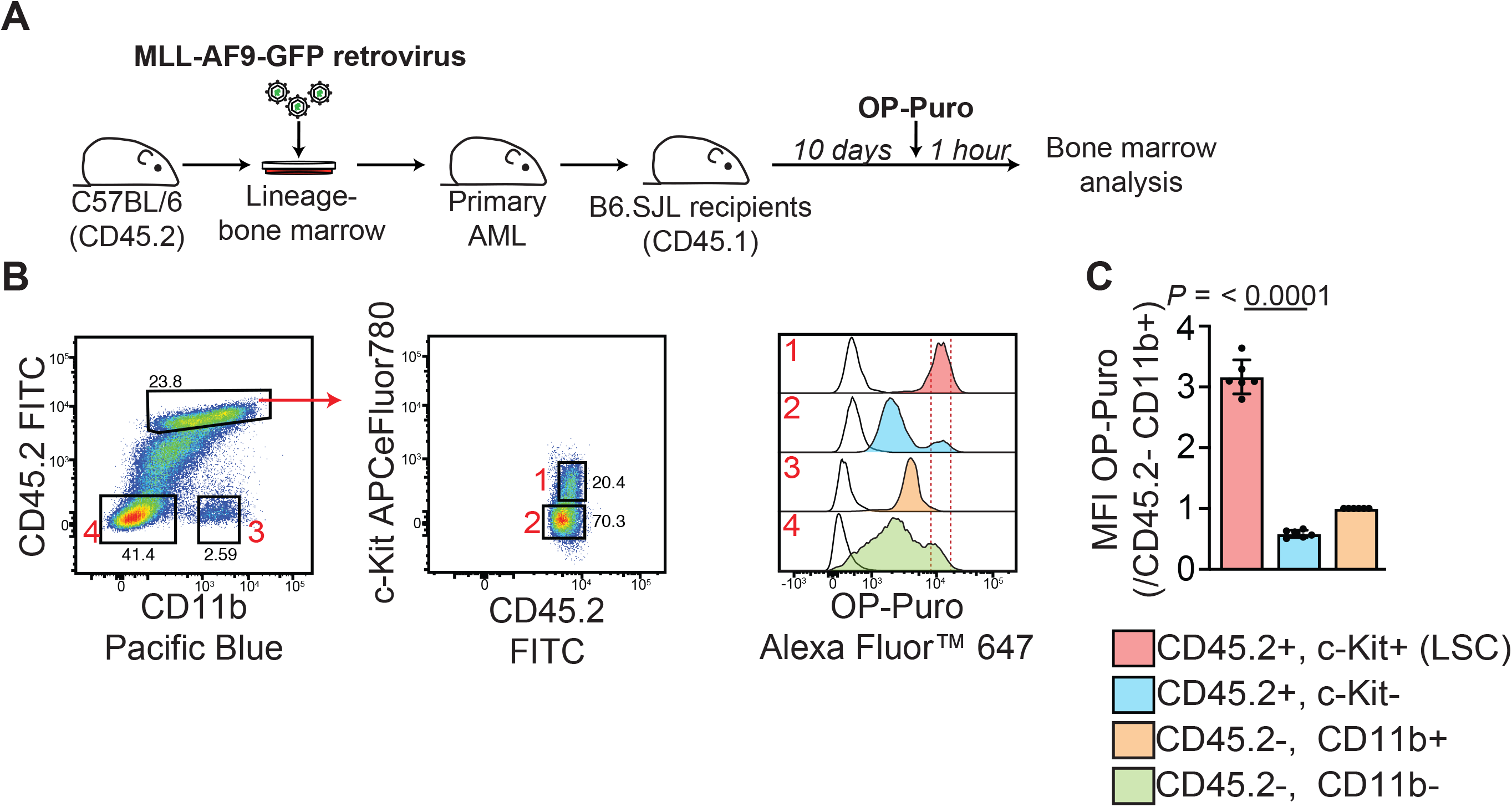
Leukemia stem cells show high protein synthesis rate. (A) Schematic overview of the O-propargyl puromycin (OP-Puro) labeling experiment. (B) Representative flow cytometry gating strategy to quantify OP-Puro incorporation in the BM of leukemia-engrafted mice. CD45.2 was used to identify leukemia cells due to the loss of GFP fluorescence during cell fixation and permeabilization. The white curves represent BM of PBS-injected control mouse. (C) Median fluorescence intensity (MFI) of OP-Puro normalized against CD45.2-/CD11b+ host myeloid precursor cells. The data represents two independent experiments with a total of six animals per group. All graphs show mean +/- standard deviation. Student’s *t*-test was used to determine statistical significance. Two-tailed *P* values are shown.

### Defective ribosome assembly significantly extends survival in leukemia-engrafted mice

To assess the significance of high protein synthesis rate on LSC function and leukemia propagation, we took advantage of a recently reported transgenic eukaryotic initiation factor 6 (eIF6) mouse strain^18^. These mice express human *EIF6* transgene by a doxycycline (Dox)-responsive promoter located downstream of the *Collagen 1a1* (*Col1a1*) gene, allowing inducible and graded overexpression of eIF6 **(Figure 2A)**. Given its ability to bind the intersubunit face of 60S ribosomal subunits, eIF6 functions as a ribosome anti-association factor that prevents the joining of 60S and 40S subunits into translation-competent 80S ribosomes^18,19^. Enforced expression of eIF6 therefore disrupts ribosomal subunit joining and attenuates global protein synthesis rate^18^.

**Figure 2.**
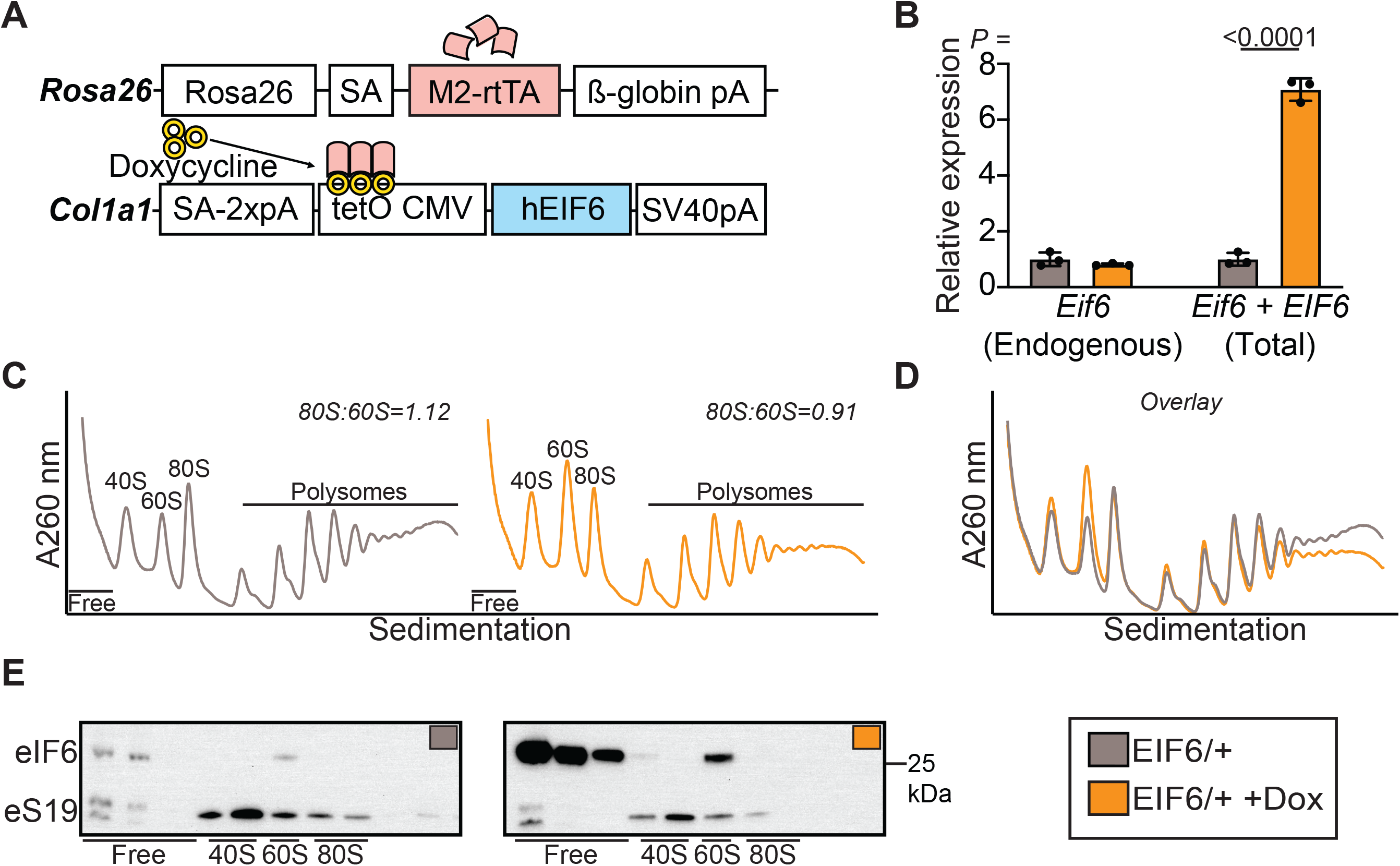
Enforced expression of eIF6 impairs ribosomal subunit joining. (A) Schematic overview of transgenic eIF6 mouse strain. *M2-reverse tetracycline transactivator* (*M2-rtTA*) is constitutively expressed at the *Rosa26* locus. This permits doxycycline-inducible expression of the human *EIF6* transgene at downstream of the *Col1a1* locus. (B) Quantitative reverse transcription PCR analysis of endogenous murine *Eif6* and total murine *Eif6* + transgenic *EIF6* mRNA in prospectively purified c-Kit+ leukemia cells. Cells were harvested from leukemia-engrafted mice that were administered Dox for five days. n=3 biologically independent samples per group. (C, D) Sucrose gradient sedimentation of extracts from cultured c-Kit+ leukemia cells. Dox induction, 16 hr. Shown is representative of two independent experiments. (E) Immunoblot analysis of eIF6 and eS19 (Rps19) in fractionated extracts. Shown is representative of two independent experiments. All graphs show mean +/- standard deviation. Student’s *t*-test was used to determine statistical significance. Two-tailed *P* values are shown.

We isolated BM cells from mice with two copies of *M2-rtTA* transgene and one copy of *EIF6* transgene and transduced them with MLL-AF9-GFP retrovirus to generate an aggressive MLL-AF9 AML where the expression of *EIF6* transgene is Dox-inducible. To evaluate the level of *EIF6* transgene expression, we isolated c-Kit+ LSCs from Dox-treated leukemia-engrafted mice and performed quantitative reverse transcription PCR (RT-qPCR) to measure *EIF6* mRNA. Using two sets of primers to distinguish endogenous mouse *Eif6* mRNA from total (endogenous + transgene) *EIF6* mRNA, we could observe a 7.1-fold increase in *EIF6* mRNA compared with controls **(Figure 2B)**. To assess the impact of eIF6 overexpression on ribosome assembly, we cultured freshly harvested c-Kit+ LSCs in the presence of Dox and profiled cell extracts by sucrose gradient sedimentation. Consistent with a ribosomal subunit joining defect, eIF6 overexpression led to increased levels of free ribosomal subunits with a concomitant shift in polysome distribution toward the lighter polysomes **(Figure 2C and 2D)**. Immunoblotting of fractionated cell extracts confirmed a robust increase in eIF6 protein abundance, both in the ribosome-free fraction and in association with 60S subunits **(Figure 2E)**.

Next, we transplanted sublethally irradiated recipient mice with leukemia cells and administered them Dox to induce eIF6 overexpression **(Figure 3A)**. Of note, Dox administration was initiated one day after transplantation to avoid confounding effects of eIF6 overexpression on leukemia cell engraftment. At two weeks after transplantation, Dox-treated mice showed on average a 65% decrease in the number of white blood cells in the peripheral blood compared with controls **(Figure 3B)**. This outcome was associated with a 30% decrease in the frequency of GFP+ leukemia cells **(Figure 3C)**. Consistent with reduced leukemia burden, eIF6 overexpression significantly prolonged survival in leukemia-engrafted mice (median survival of 15.5 and 24 days in control and Dox groups, respectively) **(Figure 3D)**.

**Figure 3.**
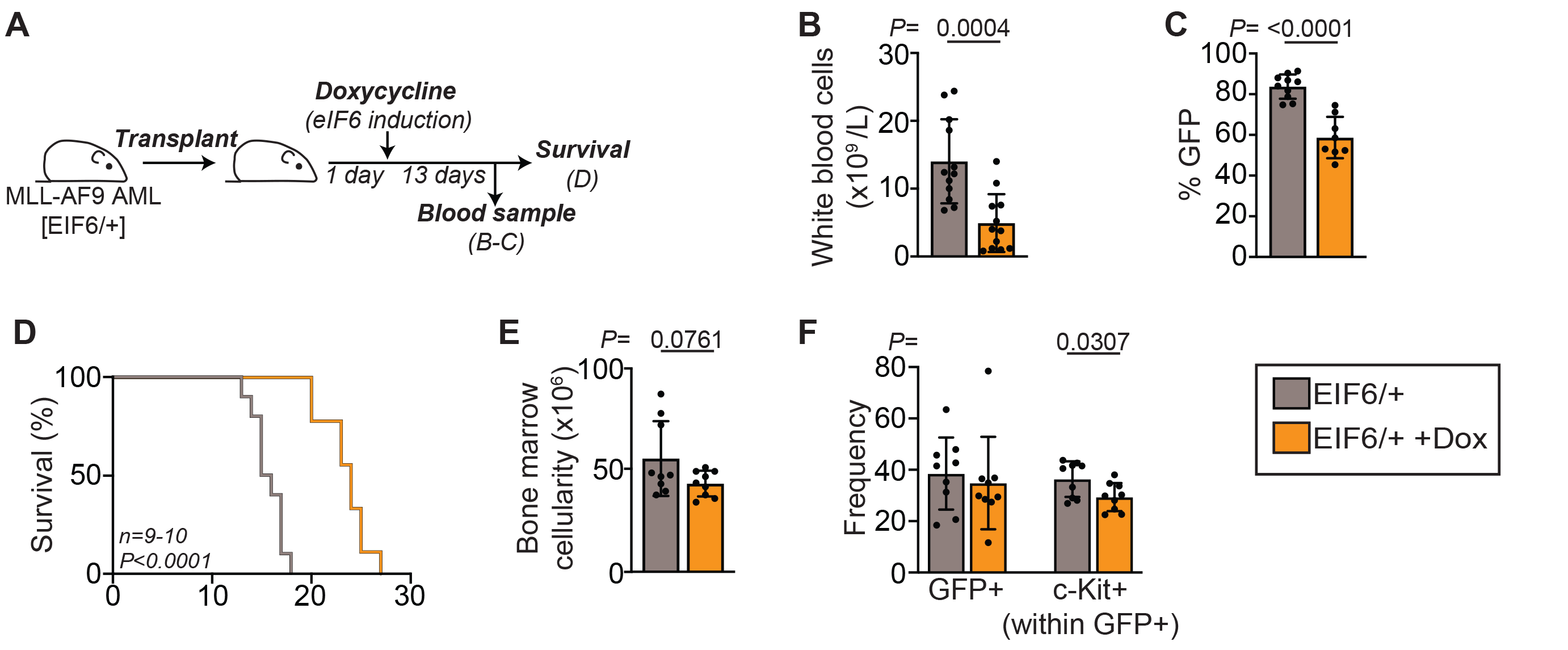
Enforced expression of eIF6 prolongs survival by eradicating leukemia stem cells. (A) Experimental strategy to assess the impact of eIF6 overexpression on leukemia propagation. 50,000 serial transplanted leukemic bone marrow cells were transplanted into sublethally irradiated recipient mice. One day after transplantation, half of the engrafted mice were administered Dox to induce *EIF6* transgene expression. (B) Number of white blood cells in peripheral blood. n=12 animals per group. (C) Frequency of GFP+ cells within the total CD45+ fraction in the peripheral blood. n=10 and 8 animals per group. (D) Survival of leukemia-engrafted mice. n=10 and 9 animals per group. (E) Bone marrow cellularity. 50,000 serial transplanted leukemic bone marrow cells were transplanted into sublethally irradiated recipient mice. Five days after transplantation, half of the engrafted mice were administered Dox and all mice were analyzed on day ten. n=9 animals per group. (F) Frequency of GFP+ leukemia cells and c-Kit+ LSCs within the GFP+ leukemia cells in the bone marrow. n=9 animals per group. All graphs show mean +/- standard deviation. Student’s *t*-test was used to determine statistical significance in B, C, E and F. Two-tailed *P* values are shown. Log-rank (Mantel Cox) test was used to determine statistical significance in D.

To assess the short-term impact of eIF6 overexpression on c-Kit+ LSCs and their progeny, we harvested BM cells from mice that were administered Dox for five days and quantified leukemia cells by flow cytometry **(Supplementary Figure 2)**. In the BM, induction of eIF6 overexpression had no significant impact on total cellularity or on the frequency of GFP+ leukemia cells **(Figure 3E and 3F)** but led to a significant reduction in the frequency of c-Kit+ LSCs within the GFP+ leukemia cell compartment **(Figure 3F)**. Spleen analysis revealed a significant decrease in spleen weight in Dox-treated mice, and a reduction in both GFP+ leukemia cell and c-Kit+ LSC fractions, although not statistically significant **(Supplementary Figure 3A-3C)**. Taken together, these data suggest that defective ribosome assembly prolongs survival by impairing LSC function.

### A severe ribosome assembly defect selectively eradicates LSCs

The significant impact of eIF6 overexpression on leukemia propagation prompted us to assess whether a more severe ribosome assembly defect was sufficient to completely eradicate the disease. To this end, we generated an aggressive MLL-AF9 AML using BM cells from mice with two copies of *M2-rtTA* and *EIF6* transgenes. The presence of an additional copy of *EIF6* transgene in this AML model increased the level of total *EIF6* mRNA overexpression to 12.2-fold compared with 7.1-fold in the model with one copy of *EIF6* transgene **(Supplementary Figure 4A, see also Figure 2B)**. Consistent with a dose-dependent effect of eIF6 on ribosomal subunit joining^18^, induction of high eIF6 overexpression led to a rapid collapse of polysomes **(Supplementary Figure 4B-4C)**.

High eIF6 overexpression in leukemia-engrafted mice further extended the median survival to 31.5 days compared with 24 days in the model with one *EIF6* transgene **(Figure 4A)**. Consistent with a more profound impact on leukemia propagation, a short-term induction of high eIF6 overexpression led to a significant reduction in BM cellularity **(Figure 4B)** that was associated with 83% decrease in the frequency of GFP+ leukemia cells **(Figure 4C)**. The frequency of c-Kit+ LSCs within the GFP+ compartment was reduced by 62% **(Figure 4C)**. Of interest, we could observe a 29% and 53% decrease in GFP and c-Kit median fluorescence intensities, respectively, in c-Kit+ LSCs compared with controls **(Figure 4D and 4E)**. No differences were observed in the ability of c-Kit+ leukemia cells to form colonies **(Figure 4F)**. Analysis of the spleens revealed a significant reduction in spleen weight that was associated with a significant decrease in GFP+ leukemia cells and c-Kit+ LSCs **(Supplementary Figure 4D-4E**).

**Figure 4.**
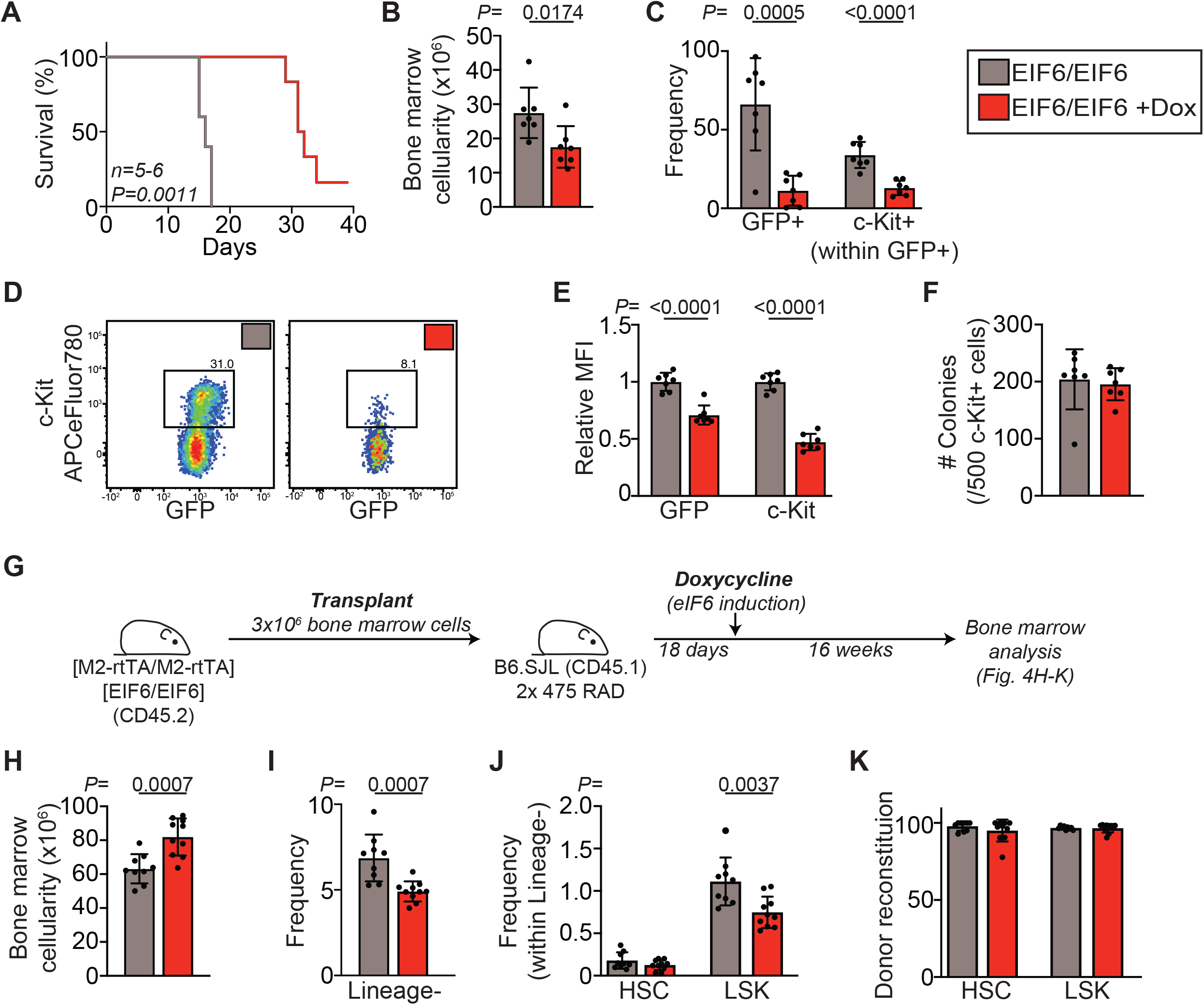
A severe ribosome assembly defect selectively eradicates LSCs. (A) Survival of leukemia-engrafted mice. 100,000 serial transplanted leukemic bone marrow cells were engrafted into sublethally irradiated recipient mice. One day after transplantation, half of the engrafted mice were administered Dox to induce *EIF6* transgene expression. n=5 and 6 animals per group. (B) Bone marrow cellularity. 100,000 serial transplanted leukemic bone marrow cells were engrafted into sublethally irradiated recipient mice. Five days after transplantation, half of the mice were administered Dox and all mice were analyzed on day ten. n=7 animals per group. (C) Frequency of GFP+ leukemia cells and c-Kit+ LSCs within the GFP+ leukemia cells in the bone marrow. n=7 animals per group. (D) Representative flow cytometry plots of c-Kit+ LSCs. (E) Median fluorescence intensity (MFI) of GFP and c-Kit within the c-Kit+ LSC compartment. n=7 animals per group. (F) Colony-forming potential of c-Kit+ LSCs. Five hundred prospectively purified c-Kit+ LSCs were isolated from Dox-treated mice and seeded on semi-solid methylcellulose. Data represents two independent experiments with a total of seven biologically independent samples per group. (G) Schematic overview of the non-competitive transplantation experiment. Three million unfractionated bone marrow cells isolated from [M2-rtTA/M2-rtTA][EIF6/EIF6] mice (CD45.2+) were transplanted into lethally irradiated recipient mice (CD45.1+). Eighteen days after transplantation, half of the recipients were administered Dox to induce eIF6 overexpression. Bone marrow analysis was performed 16 weeks after Dox administration. (H) Bone marrow cellularity. n=9 and 10 animals per group. (I) Frequency of Lineage-cells in the bone marrow. n=9 and 10 animals per group. (J) Frequency of HSCs and LSK cells within the Lineage-bone marrow cells. n=9 and 10 animals per group. (K) Donor cell reconstitution within the HSC and LSK cell compartments. n=9 and 10 animals per group. All graphs show mean +/- standard deviation. Student’s *t*-test was used to determine statistical significance. Two-tailed *P* values are shown. Log-rank (Mantel Cox) test was used to determine statistical significance in A. HSC, hematopoietic stem cell; LSK, Lineage-/Sca-1+/c-Kit+.

To compare the selectivity of leukemia cells and normal hematopoietic cells to defective ribosome assembly, we next performed a non-competitive transplantation experiment using BM cells isolated from mice with two copies of *M2-rtTA* and *EIF6* transgenes **(Figure 4G)**. BM analysis of recipient mice that were administered Dox for 16 weeks revealed a significant increase in total cellularity and a significant decrease in the frequency of lineage-negative cells **(Figure 4H and 4I, Supplementary Figure 4F)**. This suggests that the increase in BM cellularity is due to an accumulation of lineage-committed precursor cells. Further analysis of the more immature stem and progenitor cell compartment revealed a significant decrease in the frequency of Lineage-/Sca-1+/c-Kit+ (LSK) cells, with no significant difference in the frequency of hematopoietic stem cells (HSCs) **(Figure 4J)**. Importantly, both HSC and LSK compartments were devoid of host-derived cells **(Figure 4K)**. Taken together, despite the dramatic effect on LSC function and leukemia propagation, high eIF6 overexpression had only a modest effect on the frequency of normal hematopoietic stem and progenitor cells in the BM. These results indicate that LSCs are selectively sensitive to defective ribosome assembly.

### A rare subset of LSCs with immature stem cell signature underlies resistance to defective ribosome assembly

To obtain insights into the mechanisms that enable a subset of c-Kit+ LSCs to adapt to defective ribosome assembly, we first performed transcriptional profiling of c-Kit+ LSCs isolated from leukemia-engrafted mice that were administered Dox for five days. Statistical analysis of the normalized data identified a total of 800 differentially expressed genes (adjusted *P* < 0.05, at least 1.5-fold change (FC)) **(Figure 5A, Supplementary File 1)**. RT-qPCR analysis of *EIF6* and three additional genes verified the accuracy of the transcriptome dataset **(Supplementary Figure 5A-5B)**. Principal component analysis (PCA) clustered the samples by group with PC1 explaining 77% of the variation **(Figure 5B)**. To address whether differentially expressed genes reflected a shift in hematopoietic lineage identity of c-Kit+ LSCs, we compared them to a previously published dataset of normal murine hematopoiesis using CellRadar^20^. We observed that eIF6 overexpression led to upregulation of genes that associate with HSC identity and downregulation of genes that associate with the granulocyte and monocyte identities **(Figure 5C)**. These results demonstrate that the induction of defective ribosome assembly leads to enrichment of LSCs that display a relatively immature transcriptional identity.

**Figure 5.**
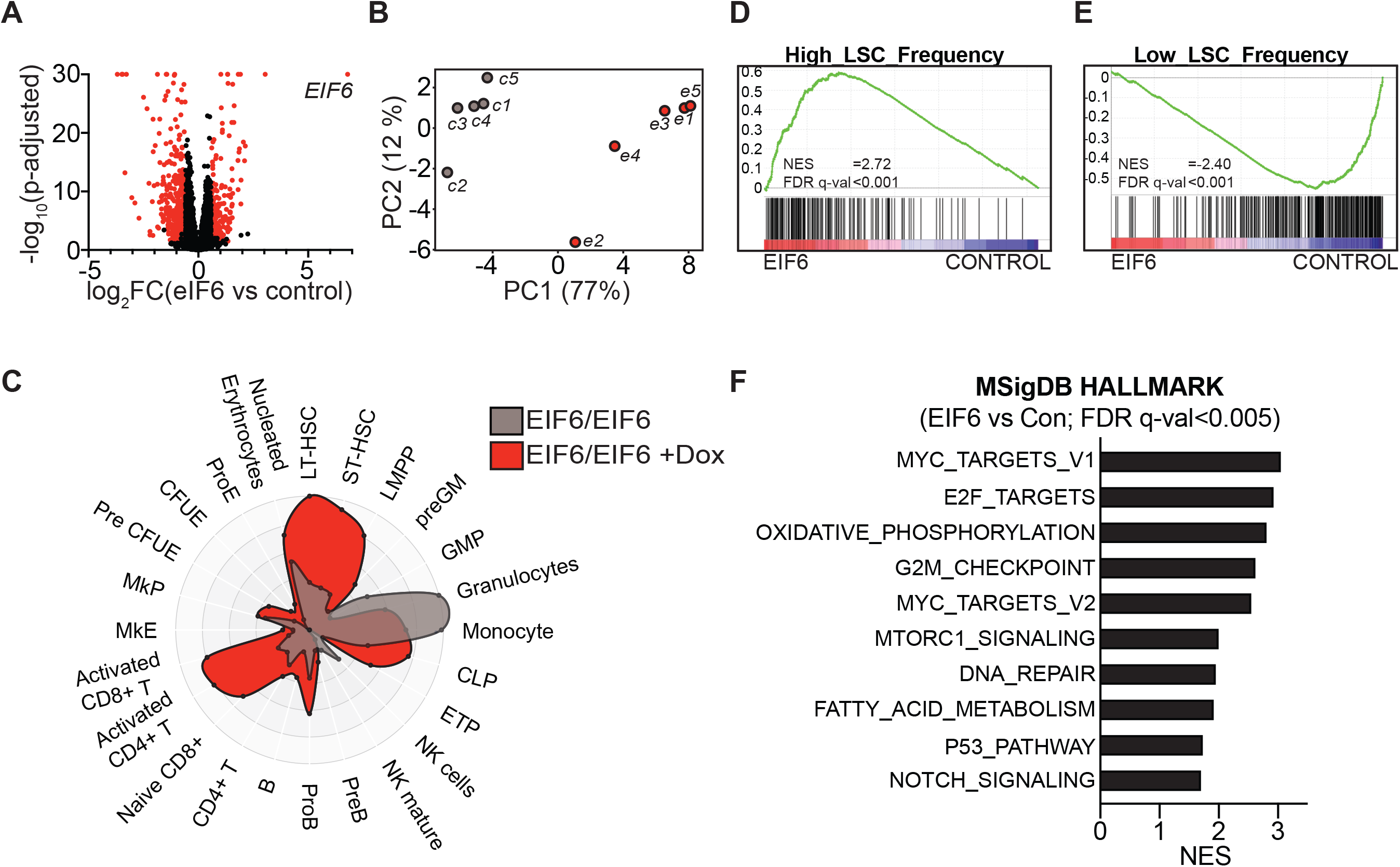
Resistance to defective ribosome assembly emerges from a rare subset of leukemia stem cells. (A) Volcano plot illustration of differential gene expression in c-Kit+ LSCs. Red dots depict differentially expressed genes (adjusted *P* < 0.05, at least 1.5-fold change). n=5 biologically independent samples per group. FC, fold change. (B) Principal component analysis of transcriptomic data. c, control; e, eIF6. (C) CellRadar plot of differentially expressed genes (adjusted *P* < 0.05, at least 1.5-fold change). (D-E) Gene Set Enrichment Analysis of transcriptomic data using previously published LSC gene sets^21^. (F) Summary of the ten most significantly enriched Molecular Signatures Database Hallmark gene sets^22^. NES, normalized enrichment score; FDR, false discovery rate.

To address the association of eIF6^high^ leukemia cells with LSC identity more directly, we next performed Gene Set Enrichment Analysis (GSEA)^13^ using previously published gene signatures that correlate with LSC frequency in mouse models of MLL-rearranged AML^21^. We observed a significant enrichment of the genes that correlate with high LSC frequency among genes upregulated upon eIF6 overexpression (**Figure 5D**). Conversely, genes that were downregulated were significantly enriched for those that correlate with low LSC frequency (**Figure 5E**). Further GSEA using the Molecular Signatures Database (MSigDB) Hallmark gene set collection^22^ revealed a significant enrichment in response to eIF6 overexpression of molecular processes that have been associated with LSC function, including Myc targets^23^, oxidative phosphorylation^24^ and fatty acid metabolism^25^ (**Figure 5F and Supplementary Table 2**). Conversely, downregulated genes were significantly enriched for Inflammatory response and Coagulation gene sets **(Supplementary Table 3)**. Taken together, these data indicate that the resistance to defective ribosome assembly emerges from a distinct subset of LSCs with an immature stem cell molecular signature.

### Resistant LSCs exhibit high ribosome content

GSEA identified a significant enrichment of genes encoding ribosomal proteins and assembly factors among those upregulated in response to eIF6 overexpression **(Figure 6A, left panel)**. To address whether the abundance of these factors was also increased at the protein level, we performed quantitative proteomics on c-Kit+ LSCs. We detected a total of 8517 proteins, of which 189 were differentially expressed in response to eIF6 overexpression (adjusted *P* < 0.05, at least 1.5-FC) (**Supplementary Figure 6A, Supplementary File 2**). These included eIF6, which showed a 7.9-fold increase compared with controls, as well as GFP and c-Kit that were decreased by 44% and 29%, respectively **(Supplementary File 2)**, consistent with flow cytometry data (**Figure 4D-4E**). The proteome and transcriptome datasets showed a significant correlation (Pearson correlation coefficient r=0.56) with only four proteins showing strong upregulation at protein level while the corresponding transcript was unchanged **(Supplementary Figure 6B)**. GSEA of the two datasets identified overlapping Hallmark gene sets **(Supplementary Figure 6C)**, and the proteome dataset confirmed a significant enrichment of ribosomal proteins and assembly factors upon eIF6 overexpression **(Figure 6A, right panel)**. Mitochondrial ribosomal proteins were also enriched both at mRNA and protein level **(Figure 6B)**. Of interest, the 60S subunit assembly factors Znf622, Nmd3 and Rsl24d1 were identified among the eight most strongly higher expressed ribosomal factors, while their respective mRNAs showed no significant change compared with controls **(Figure 6C, Supplementary Figure 6D)**. Since these factors are evicted from the nascent pre-60S subunit in the cytoplasm, this finding suggests that eIF6 overexpression may interfere with the final maturation of pre-60S subunits, leading to accumulation of such particles^26^.

**Figure 6.**
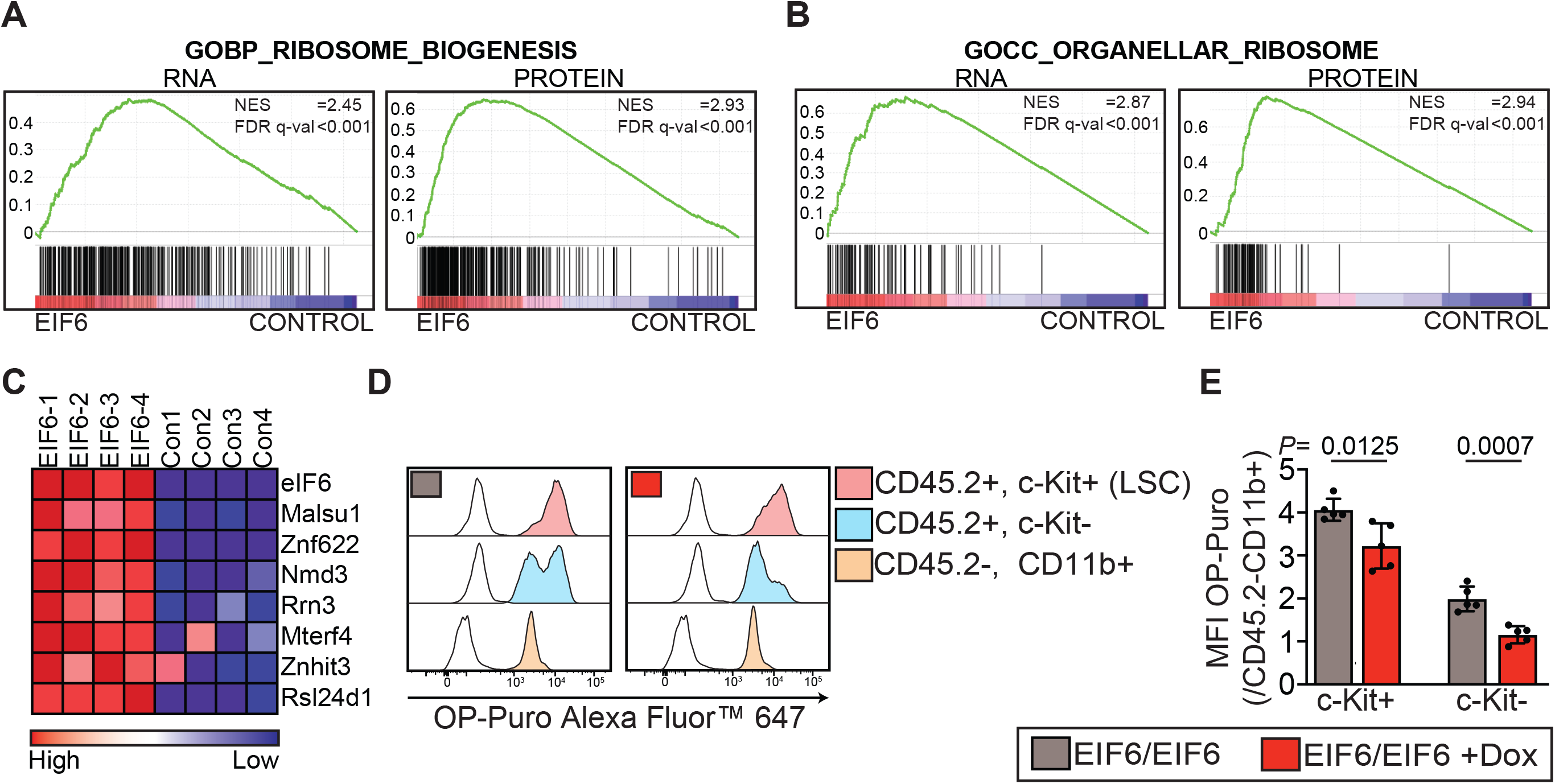
Resistant LSCs exhibit high ribosome content. (A) Gene Set Enrichment Analysis of the transcriptome and proteome datasets using ribosome biogenesis gene signature. (B) Gene Set Enrichment Analysis of the transcriptome and proteome datasets using organellar ribosome gene signature. (C) Heatmap of the eight most highly expressed ribosomal proteins and assembly factors shown in (A, right panel) in response to eIF6 overexpression. Data is represented as row-normalized expression values. (D) Representative flow cytometry plot of OP-Puro incorporation in the bone marrow of leukemia-engrafted mice. CD45.2 was used to identify leukemia cells due to the loss GFP fluorescence during cell fixation and permeabilization. The white curves represent bone marrow of PBS-injected control mouse. (E) Median fluorescence intensity (MFI) of OP-Puro normalized against the CD45.2-/CD11b+ host myeloid precursor cells. The data represents two independent experiments with a total of five animals per group. All graphs show mean +/- standard deviation. Student’s *t*-test was used to determine statistical significance. Two-tailed *P* values are shown. NES, normalized enrichment score; FDR, false discovery rate.

To address whether the persistent LSCs with high ribosome content continued to show attenuated global protein synthesis rate, we next performed OP-Puro labeling experiments in leukemia-engrafted recipient mice. Consistent with our earlier analysis **(Figure 1)**, we observed that c-Kit+ LSCs in control mice incorporated approximately four times more OP-Puro compared with normal host-derived CD11b+ myeloid BM cells, and that c-Kit- leukemia cells could be divided into OP-Puro^high^ and OP-Puro^low^ subsets **(Figure 6D)**. The induction of high eIF6 overexpression led to a 21% and 42% decrease in OP-puro incorporation in c-Kit+ and c-Kit- leukemia cells, respectively **(Figure 6E)**. The more pronounced reduction in OP-Puro incorporation in c-Kit- leukemia cells was associated with a selective loss of the OP-Puro^high^ cell subset.

### Loss of p53 does not alter the impact of defective ribosome assembly on leukemia propagation

The transcriptome and proteome datasets revealed a positive enrichment of the p53 pathway signature in response to eIF6 overexpression, including a 1.8-fold increase in p53 protein (**Figure 7A and 7B)**. This finding is of interest as perturbations of ribosome biogenesis are known to trigger a p53-dependent ribosome biogenesis stress pathway^27,28^. To assess the role of p53 in the depletion of LSCs upon defective ribosome assembly, we applied CRISPR/Cas9 to generate a p53-deficient AML subclone **(Supplementary Figure 7A)**. We first cultured p53-proficient and p53-deficient leukemia cells in the presence of Dox, and measured cell viability using trypan blue exclusion assay. This experiment revealed a robust induction of apoptosis in response to eIF6 overexpression regardless of p53 status **(Supplementary Figure 7B)**. By contrast, treatment of these cells with the Mdm2 inhibitor Nutlin-3a or Actinomycin D, a known inducer of ribosome biogenesis stress pathway^28^, induced apoptosis in p53-proficient but not in p53-deficient cells (**Supplementary Figure 7B**). Consistent with these results, loss of p53 did not alter the impact of eIF6 overexpression on leukemia propagation in vivo compared with p53-proficient parental cells (**Figure 7C, see also Figure 4A**). BM analysis of mice transplanted with these cells revealed no difference in total BM cellularity after five days of Dox administration **(Figure 7D)**. However, similarly as observed with p53-proficient cells **(Figure 4C)**, we observed a 61% reduction in GFP+ leukemia cells that was associated with a 39% reduction in c-Kit+ cells within this fraction (**Figure 7E)**. Taken together, these results demonstrate that the impact of defective ribosome assembly on leukemia propagation is predominantly independent of p53 pathway.

**Figure 7.**
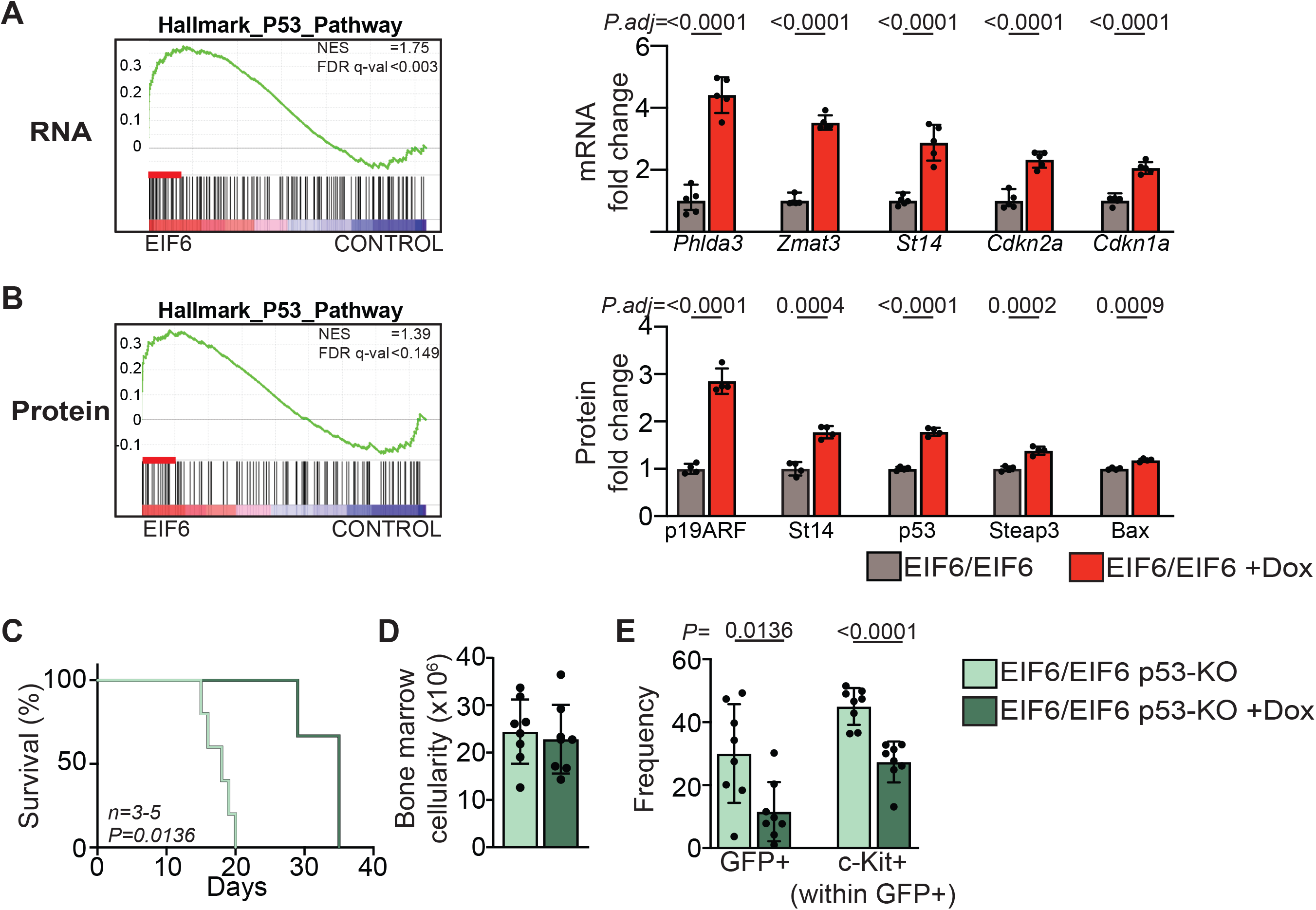
Loss of p53 does not alter the impact of defective ribosome assembly on leukemia latency. (A-B) Visualization of selected mRNAs and proteins from the leading edge subset of MSigDB Hallmark p53 pathway gene set^22^. Leading edge (depicted as a red bar) represents the genes that contribute the most to the enrichment signal of a given gene signature. (C) Survival of leukemia-engrafted mice. 100,000 serial transplanted leukemic bone marrow cells were engrafted into sublethally irradiated recipient mice. Doxycycline administration was initiated one day after transplantation. n=5 and 3 animals per group. (D) Bone marrow cellularity. 100,000 serial transplanted leukemic bone marrow cells were engrafted into sublethally irradiated recipient mice. Five days after transplantation, half of the mice were administered Dox and all mice were analyzed on day ten. The data represents two independent experiments with a total of eight animals per group. (E) Frequency of GFP+ leukemia cells and c-Kit+ LSCs within the GFP+ leukemia cells in the bone marrow. The data represents two independent experiments with a total of eight animals per group. All graphs show mean +/- standard deviation. Adjusted *P* values from transcriptomic and proteomic analyses are shown in A and B. Student’s *t*-test was used to determine statistical significance in D and E. Two-tailed *P* values are shown. Log-rank (Mantel Cox) test was used to determine statistical significance in C. NES, normalized enrichment score; FDR, false discovery rate.

## DISCUSSION

Mutations in genes that encode components of oncogenic signal transduction pathways represent the most common class of genetic aberrations in AML and converge on protein synthesis machinery to regulate its function^2,8^. Moreover, the transcription factor Meis1 was recently shown to support leukemogenesis by inducing transcription of ribosomal genes, suggesting a direct role for increased ribosome biogenesis in the establishment and propagation of LSCs^29^. In line with these findings, we show that LSCs in MLL-rearranged AML are characterized by high global protein synthesis rate and that they are selectively sensitive to defective ribosome assembly compared with normal hematopoietic stem and progenitor cells. Despite the significant enrichment of LSCs within c-Kit+ leukemia cells, c-Kit- negative leukemia cells are also known to contain residual LSC activity^16^. We could observe that a fraction of c-Kit- leukemia cells displayed a high protein synthesis rate comparable to that observed in c-Kit+ LSCs, and that these cells were selectively eliminated upon eIF6 overexpression. Future studies will be needed to assess the association between high protein synthesis rate and LSC activity within c-Kit- cell subset.

Despite the dramatic reduction in leukemia cells in the BM, induction of defective ribosome assembly was not sufficient to eradicate the disease. This outcome was associated with an enrichment of a distinct subset of c-Kit+ leukemia cells, suggesting that the resistance to defective ribosome assembly emerges from these cells. Molecular profiling of these cells revealed a significant enrichment for LSC gene signatures, even though these experiments were performed on c-Kit+ leukemia cells that per se are already enriched in LSCs^16^. Strikingly, these cells also exhibited a significant increase in the abundance of ribosomal proteins and assembly factors, suggesting that the ability of LSCs to adapt to defective ribosome assembly may depend on maintaining a critical threshold level of protein synthesis. Consistent with this concept, the impaired ribosome biogenesis rate in AML with *RUNX1* mutations was recently shown to confer increased sensitivity of leukemia cells to the protein synthesis inhibitor homoharringtonine^30^. Whether the observed increase in ribosome content in resistant LSCs represents an adaptative response or rather a selection for a specific LSC subtype remains to be established.

Perturbations of ribosome biogenesis in the nucleolus are known to activate a p53-dependent ribosome biogenesis stress pathway by redirecting nascent 5S ribonucleoprotein particle (5S RNP; composed of uL5, uL18 and 5S rRNA) from the assembly into pre-60S ribosomal subunit toward inhibition of Mdm2, a key negative regulator of p53^27,28^. We have previously shown that induction of the 5S RNP-Mdm2-p53 pathway leads to a transient reduction in overall leukemia burden but fails to prolong survival in murine models of MLL-rearranged AML^31^. By contrast, our present study demonstrates that perturbation of cytoplasmic ribosome assembly eradicates LSCs and significantly extends survival in leukemia-engrafted mice. Importantly, the observed outcome was not altered by loss of p53, demonstrating that the impact of defective ribosome assembly on leukemia propagation is predominantly independent of p53. These data are consistent with a recent identification of the cytoplasmic 40S ribosomal subunit assembly factor RIO-kinase 2 as a potential therapeutic target in AML regardless of p53 status^32^.

In summary, our study establishes ribosome assembly as a critical vulnerability in AML, suggesting that strategies that perturb ribosome homeostasis in the cytoplasm may find therapeutic utility in AML. Future work will be required to assess whether the adaptation of LSCs to defective ribosome assembly associates with an acquisition of novel molecular vulnerabilities that can be cotargeted to increase the therapeutic efficacy of protein synthesis inhibitors in AML.

## ACKNOWLEDGEMENTS

We thank Anna Hogmalm for technical assistance. This work was supported by the Swedish Childhood Cancer Fund (TJ2018-0022, PR2018-0022, PR2021-0015, TJ2021-0009), The Jeansson Foundation (S2018-0030), The Assar Gabrielsson Foundation (FB19-34, FB20-38, FB21-37), The Åke Wiberg Foundation (M20-0181), The Swedish Research Council (2019-01761), The Johannes and Sonja Magnusson Foundation, The Martin Odin Foundation and Sahlgrenska Academy. The authors acknowledge support from the National Genomics Infrastructure in Stockholm funded by Science for Life Laboratory, the Knut and Alice Wallenberg Foundation and the Swedish Research Council, and SNIC/Uppsala Multidisciplinary Center for Advanced Computational Science for assistance with massively parallel sequencing and access to the UPPMAX computational infrastructure.

## AUTHORSHIP CONTRIBUTIONS

P.J. conceived the study. D.S., C.G. and P.J. designed, performed and analyzed experiments. S.G. and J.H. performed mass spectrometry and protein quantification. P.J. wrote the manuscript with feedback from all authors.

## DISCLOSURE OF CONFLICT OF INTEREST

The authors declare no conflicting interests.

## Notes

### Competing Interest Statement

The authors have declared no competing interest.

