## Supplemental information for "Defective ribosome assembly impairs leukemia stem cell function in a murine model of acute myeloid leukemia"

#### Supplementary Material and Methods

##### Animal studies

Mice were housed in individually ventilated cages at a temperature of 23°C (+/- 1°C) and a relative humidity of 45-70%. C57BL6J (#000664) and B6 CD45.1 (#002014) mice were purchased from the Jackson Laboratory. Transgenic eIF6 mouse strain has been described previously[18]. All experiments were performed using 8-12 weeks old young adult mice. Both male and female mice were used as recipients in transplantation studies. The gender of donor mice is described in 'Transplantation assays' section. eIF6 overexpression was induced in vivo by feeding mice with doxycycline food pellets (2000 mg/kg; Ssniff spezialdiäten GmbH).

##### Retrovirus production

The MLL-AF9-GFP retroviral vector was obtained from Addgene (#71443, pMIG-FLAG-MLL-AF9, a gift from Daisuke Nakada). Lipofectamine LTX reagent (ThermoFisher Scientific) was used to transiently transfect ecotropic Platinum-E retroviral packaging cell line (Cell Biolabs). Retroviral supernatant was harvested 48 hours after transfection, passed through a 0.45 μm filter and stored at -80 °C.

**Retroviral transduction**

Freshly harvested bone marrow cells were stained with biotin-conjugated lineage antibodies **(Supplementary Table 1)**, and Anti-Biotin MicroBeads (Miltenyi) were used for lineage depletion. The lineage-depleted cells were cultured in OptiMEM I reduced Serum Media (Thermo Fisher Scientific), supplemented with 10 % fetal calf serum (FCS; Merck), penicillin/streptomycin (P/S, Life Technologies), 25 ng/mL murine stem cell factor (mSCF; PeproTech), 10 ng/mL murine interleukin 3 (mIL-3; PeproTech) and 50 ng/mL mouse thrombopoietin (PeproTech) for 24 hours. Retroviral transduction was performed using retronectin-coated plates (Takara) according to manufacturer's instructions.

**Peripheral blood**

Peripheral blood was collected from the tail vein into Microvette tubes (Sarstedt) and cellularity was analyzed using Sysmex KX-21 or Sysmex XP-300 hematology analyzers. For flow cytometry analyses, Dextran sedimentation (2 % in PBS, Merck) and ACK lysis buffer (Thermo Fisher Scientific) were applied to remove erythrocytes before antibody labeling.

**Immunoblot**

Proteins in 1x NuPAGE LDS sample buffer supplemented with 50 mM DTT were incubated at 85 °C for 10 min and run on NuPAGE Bis-Tris polyacrylamide gels in NuPAGE MOPS SDS running buffer (Thermo Fisher Scientific). The iBlot 2 gel transfer device (Thermo Fisher Scientific) was used to transfer proteins to nitrocellulose membranes. Membranes were first blocked for one hour in 5 % milk (Semper) in PBS supplemented with 0.1% Tween (Merck), and then stained with primary antibodies overnight at 4 °C on a tube roller. The blots were washed three times with PBS-tween and incubated for one hour with horseradish peroxidase-conjugated secondary antibody. Following five washes with PBS-tween, the blots were detected using the SuperSignal West Pico PLUS reagents (Thermo Fisher Scientific) and Fujifilm LAS-1000 Intelligent Dark Box II. Antibodies are listed in the Supplementary Table 1.

**Quantitative reverse transcription PCR**

Total RNA was isolated using RNeasy mini kit (Qiagen) and reverse transcription was performed using SuperScript III reverse trancriptase (Thermo Fisher Scientific). Real-time PCR was performed using SsoFast EvaGreen supermix (Bio-Rad) on Applied Biosystems 7900HT Real Time PCR System. Primers are listed in the Supplementary Table 4.

**Cell counting**

Unless stated otherwise, cells were counted on TC20™ Automated Cell Counter (Bio-Rad) using trypan blue. A cell size gate of ≥6 µm was applied to exclude red blood cells when counting bone marrow cellularity.

**CRISPR/Cas9 targeting**

To inactivate the *Trp53* gene, ribonucleoprotein complexes containing crRNA (target sequence: 5'-CTTCCACCCGGATAAGATGC-3'), tracrRNA and recombinant Cas9 nuclease were assembled according to manufacturer's (IDT) guidelines and delivered into leukemia cells using Human CD34+ Cell Nucleofector™ Kit and Nucleofector 2b Device with program U-008 (Lonza). After 48 hours, the transfected cells were treated with 10 µM Nutlin-3a to select for p53-deficient clones, and individual subclones were selected by plating these cells on semi-solid media (MethoCult M3231, StemCell technologies) supplied with 50 ng/mL mSCF and 10 ng/mL mIL-3. A similar strategy was applied to inactivate the two *pac* puromycin resistance genes in [M2-rtTA/M2-rtTA][EIF6/EIF6] leukemia cells (target sequence: 5'-CCACCCGCGACGACGTCCCC-3'), followed by in vitro selection of colonies using puromycin (1µg/mL, InvivoGen). Sanger sequencing was used to verify biallelic frameshift mutations in the *pac* gene (data not shown). These cells were used in OP-Puro labeling experiments shown in Figure 6D and 6E.

**Polysome profiling**

Freshly harvested bone marrow cells were stained with biotin-conjugated c-Kit antibody and enriched using Anti-Biotin MicroBeads (Miltenyi). The c-Kit-enriched cells were cultured for 16 hours in OptiMEM I media supplemented with 10 % FCS, P/S, 25 ng/mL mSCF, 10 ng/mL mIL-3 and 1µg/mL Dox, and then treated with cycloheximide (CHX; 100 µg/mL; Merck) for 5 min at 37 °C before harvesting by centrifugation. Cells were washed twice with ice-cold PBS supplemented with CHX (100 µg/mL), and lysed for 30 min on ice in lysis buffer (20 mM Hepes pH 7.5, 50 mM KCl, 10 mM $Mg(CH_3COO)_2$, 1.5 mM Dithiothreitol (DTT; Merck), 100 µg/mL CHX, cOmplete™ EDTA-free Protease Inhibitor Cocktail (Merck), RNaseOUT recombinant ribonuclease inhibitor (200 U/mL; Thermo Fisher Scientific) and IGEPAL® CA-630 (0.7 %; Merck)). The lysate was cleared by centrifugation (14,000g for 5 min at 4 °C), and RNA content was quantified using NanoDrop spectrophotometer (Thermo Fisher Scientific). Equal $A_{260}$ units (typically 3-5 units) of lysate were loaded onto a 10-50 % (w/v) sucrose gradient (prepared in 20 mM Hepes pH 7.5, 50 mM KCl, 10 mM $Mg(CH_3COO)_2$, 1.5 mM DTT, 100 µg/mL CHX, cOmplete™ EDTA-free Protease Inhibitor Cocktail, RNaseOUT recombinant ribonuclease inhibitor (4 U/mL)) prepared in open-top polyclear tubes (#7030; Seton Scientific). A Gradient Master (Biocomp) was used to prepare the sucrose gradients. After centrifugation at 36,000 rpm for 2 hours at 4 °C using a Thermo Scientific™ TH-641 rotor, sucrose gradients were profiled and fractionated using Biocomp gradient station. Proteins were precipitated with trichloroacetic acid (Merck; 25 % (vol/vol); 15 min on ice). Following centrifugation at 14,000 g for 5 min at 4 °C, protein precipitates were washed with ice-cold acetone. After drying, protein pellets were resuspended in 1x NuPAGE LDS Sample Buffer (Thermo Fisher Scientific).

**Sample preparation for mass spectrometry analysis**

300,000 FACS-sorted cells were lysed in 0.1% RapiGest (Waters) in 200 mM HEPES. Cysteine disulfide bonds were reduced with 5 mM DTT for 30 min at 56°C, alkylated with 10 mM iodoacetamide for 30 min at room temperature in the dark, and proteins were digested with sequencing grade modified trypsin (enzyme:protein ratio 1:50, Promega) at 37°C overnight. Digested peptides were labeled using TMT10plex reagents (Thermo Scientific) according to the manufacturer's protocol. Briefly, vials containing 0.8 mg of TMT label were dissolved in 41 µL of anhydrous acetonitrile (ACN). Labeling was performed by addition of 5 µL dissolved TMT10plex reagrents to each sample and incubating for 30 minutes at room temperature in a thermomixer (Eppendorf) under constant shaking at 400 rpm, followed by another 5 µL TMT reagent and incubating for 30 minutes. The reaction was stopped by adding trifluoroacetic acid to a final concentration of 0.5% (v/v), and RapiGest was precipitated by further incubation at 37°C for 45 min. Following centrifugation, the supernatants were collected, and the labeled peptides were pooled, desalted with SepPak C18 cartridges (Waters), and dried by vacuum centrifugation. High-pH reverse phase (HpH-RP) fractionation was carried out as previously described[33]. Fractionated samples were dried by vacuum centrifugation and stored at −80°C until MS analysis. Prior to LC-MS analysis, samples were dissolved in 4% ACN/0.1% formic acid (FA).

**Mass spectrometry data acquisition**

TMT-labeled peptide fractions were loaded onto a trap cartridge (Thermo Fisher Scientific Acclaim PepMap C18, 5 µm, 300 µm x 5 mm) followed by EASY-Spray PepMap Neo analytical column, 2 µm, 75 µm x 50 cm (Thermo Fisher Scientific). Each fraction was injected twice and eluted over 180 min gradients (UltiMate 3000 UHPLC, Thermo Fisher Scientific) ranging from 4-19% Solvent B (0.1%FA in 80% ACN) over 83 min, 19-29% B over 40 min, 29-41% B over 8 min, 41-95% B over 2 min and held at 95% B for 8 min at a constant flow rate of 300 nl/min at 450C. MS/MS were acquired on a Thermo Orbitrap Exploris 480 equipped with FAIMS Pro interface (Thermo Fisher Scientific) in data-dependent acquisition. FAIMS compensation voltages (CV) were set to -45, -60, and -80 with a cycle time of 1s per FAIMS experiment. The spray voltage was set at 2.1 kV and the ion transfer tube temperature was set at 275 °C. The full MS scan was performed in the Orbitrap in the range of 400 to 1400 m/z at a resolution of 60,000 with automatic gain control (AGC) set to standard and maximum ion injection time set to auto. The intensity threshold was set to 5.0e4 and mass tolerance to 10 ppm. The most intense ions selected in the first MS scan were isolated for higher-energy collision-induced dissociation (HCD) at a precursor isolation window width of 0.7 m/z, and normalized AGC of 200%. Maximum ion injection time for MS2 was set to auto with MS2 resolution set to 45,000 FWHM. The first mass and the normalized collision energy were set to 110 m/z and 35%, respectively.

**Protein quantification and statistical analysis**

MS raw files were searched in Proteome Discoverer (version 2.5; Thermo Scientific) against the Swissprot mouse database (downloaded June 2021) together with human eIF6 protein sequence, GFP, MLL-AF9, as well as commonly observed contaminants using the Sequest HT node. The enzyme was set to trypsin with up to two missed cleavages, Cysteine carbamidomethylation and TMT6plex (+ 229.163 Da) were set as static modifications whereas methionine oxidation, N-terminal acetylation, and Met loss (protein N-terminal Met) were set as dynamic modifications. TMT batch-specific isotopic correction factor was applied in the reporter ion quantification. MS1 mass tolerance was set to 10 ppm and MS2 to 0.02 Da. The false discovery rate for peptide-spectrum matches (PSMs) was set to 0.01 using the Percolator node. The co-isolation threshold was set to 50. Statistical analysis of the TMT quantification was performed using MSStatsTMT package (2.4.1)[34] in R. PSMs results from Proteome Discoverer were exported and converted into MSstatsTMT-compatible format using 'PDtoMSstatsTMTFormat' function. PSMs were filtered with peptide percolator q value < 0.01. Only unique peptides were used for protein quantifications. Protein summarization was performed using the 'msstats' method and global median normalization was performed. Differential expression analysis was performed using moderated t-tests with Benjamini–Hochberg multiple hypothesis correction.

**Data analysis**

Graphs were prepared using Prism 9 (Graphpad) and Photoshop CS5 (Adobe). Figures were prepared using Illustrator CS6 (Adobe).

**Supplementary Figure Legends**

**Supplementary Figure 1. c-Kit receptor permits enrichment of leukemia stem cells in MLL-AF9 AML.**

(A) In vitro expansion of leukemia cell subsets. Cultures were seeded with 7500 prospectively purified leukemia cells and counted on day three. The data represents a single experiment with three technical replicates.

(B) Colony-forming potential of leukemia cell subsets. Five hundred prospectively purified leukemia cells were seeded on semi-solid methylcellulose and colonies were counted on day five. The data represents a single experiment with two technical replicates.

All graphs show mean +/- standard deviation. Student's *t*-test was used to determine statistical significance. Two-tailed *P* values are shown.

**Supplementary Figure 2. Experimental strategy to quantify leukemia cell subsets in the bone marrow.**

**Supplementary Figure 3. Impact of eIF6 overexpression on leukemia propagation in the spleen.**

(A) Experimental strategy to assess the impact of eIF6 overexpression on the spleen in leukemia-engrafted mice. 50,000 serial transplanted leukemic bone marrow cells were transplanted into sublethally irradiated recipient mice. Five days after transplantation, half of the mice were administered Dox and all mice were analyzed on day ten.

(B) Spleen weight. n=9 animals per group.

(C) Frequency of GFP+ leukemia cells and c-Kit+ LSCs within the GFP+ leukemia cells in the spleen. n=9 animals per group.

All graphs show mean +/- standard deviation. Student's *t*-test was used to determine statistical significance. Two-tailed *P* values are shown.

**Supplementary Figure 4. High eIF6 overexpression results in severe ribosomal subunit joining defect.**

(A) Quantitative reverse-transcription PCR analysis of total murine *Eif6* + transgenic *EIF6* mRNA in prospectively purified c-Kit+ leukemia cells. Cells were harvested from leukemia-engrafted mice that were administered Dox for five days. n=8 biologically independent samples per group.

(B-C) Sucrose gradient sedimentation of extracts from cultured leukemia cells. Cells were treated with Dox for either (B) 12 hours or (C) 16 hours. The two time-points represent independent experiments.

(D) Spleen weight. 100,000 serial transplanted leukemic bone marrow cells were engrafted into sublethally irradiated recipient mice. Five days after transplantation, half of the mice were administered Dox and spleen analysis was performed on day ten. n=4 animals per group.

(E) Frequency of GFP+ leukemia cells and c-Kit+ LSCs within the GFP+ leukemia cells in the spleen. n=4 animals per group.

(F) Flow cytometry strategy to quantify HSCs and LSK cells in the bone marrow.

All graphs show mean +/- standard deviation. Student's *t*-test was used to determine statistical significance. Two-tailed *P* values are shown.

HSC, hematopoietic stem cell; LSK, Lineage-/Sca-1+/c-Kit+.

**Supplementary Figure 5. RT-qPCR verification of RNA sequencing data.**

(A) Comparison of total murine *Eif6* + transgenic *EIF6* mRNA expression between RNA sequencing and RT-qPCR. c-Kit+ leukemia cells were purified from leukemia-engrafted mice that were treated with Dox for five days. n=5 and n=8 biologically independent samples for RNA sequencing and RT-qPCR groups, respectively. The RT-qPCR data is the same as shown in Supplementary Figure 4A.

(B) Quantification of *Rps19*, *Cybb* and *S100a9* mRNA using RNA sequencing and RT-qPCR. *Rps19* and *S100a9*: n=5 and n=4 biologically independent samples for RNA sequencing and RT-qPCR groups, respectively. *Cybb*: n=5 and n=8 biologically independent samples for RNA sequencing and RT-qPCR groups, respectively.

All graphs show mean +/- standard deviation. Student's *t*-test was used to determine statistical significance. Two-tailed *P* values are shown.

**Supplementary Figure 6. Quantitative proteomic analysis of c-Kit+ leukemia cells.**

(A) Volcano plot illustration of differential protein abundance in c-Kit+ LSCs. Red dots depict differentially abundant proteins (adjusted $P < 0.05$, at least 1.5-fold change). n=4 biologically independent samples per group. FC, fold change.

(B) Correlation between the transcriptome and proteome datasets. The Pearson correlation coefficient (r) and two tailed *P* value is shown. Highlighted are the four proteins that show strong upregulation at protein level but not at RNA level.

(C) Summary of the positively enriched (FDR q-val<0.15) Molecular Signatures Database Hallmark gene sets[22]. NES, normalized enrichment score; FDR, false discovery rate.

(D) Comparison of Znf622, Nmd3 and Rsl24d1 expression between the transcriptome and proteome datasets. Adjusted *P* values from transcriptomic and proteomic analyses are shown.

**Supplementary Figure 7. Genetic and functional characterization of the p53-deficient leukemia subclone.**

(A) Verification of biallelic deletions at the Cas9 cut site using Sanger sequencing.

(B) Cell viability. Leukemia cells were cultured in the presence of Dox, Actinomycin D (5 nM) or Nutlin-3a (10 μM) for 24 hours, and cell viability was measured using trypan blue exclusion assay. n=3 independent experiments.

All graphs show mean +/- standard deviation. Student's *t*-test was used to determine statistical significance. Two-tailed *P* values are shown.

**Supplementary Table 1. A list of antibodies used in this study.**

| Antigen | Application | Conjugate | Clone | Cat# | Manufacturer | Dilution |
|---|---|---|---|---|---|---|
| CD11b | Flow cytometry | BV421 | M1/70 | 101235 | Biolegend | 1:200 |
| c-Kit | Flow cytometry | APC-eFluor780 | 2B8 | 47-1171-82 | Thermo Fisher Scientific | 1:100 |
| Sca1 | Flow cytometry | Pacific Blue | E13-161.7 | 1212600 | Sony Biotechnology | 1:100 |
| CD48 | Flow cytometry | PE | HM48-1 | 1117025 | Sony Biotechnology | 1:200 |
| CD150 | Flow cytometry | APC | TC15-12F12.2 | 1179550 | Sony Biotechnology | 1:200 |
| CD3ε | Flow cytometry | PE/Cy5 | 145-2C11 | 1101550 | Sony Biotechnology | 1:400 |
| CD11b - Mac1 | Flow cytometry | PE/Cy5 | M1/70 | 1106050 | Sony Biotechnology | 1:400 |
| CD45R-B220 | Flow cytometry | PE/Cy5 | RA3-6B2 | 1116050 | Sony Biotechnology | 1:200 |
| Ly-6G/Ly-6C (Gr-1) | Flow cytometry | PE/Cy5 | RB6-8C5 | 1142050 | Sony Biotechnology | 1:400 |
| TER-119 | Flow cytometry | PE/Cy5 | TER-119 | 1181050 | Sony Biotechnology | 1:400 |
| CD45.1 | Flow cytometry | BUV395 | A20 | 565212 | BD Biosciences | 1:200 |
| CD45.2 | Flow cytometry | BUV737 | 104 | 612778 | BD Biosciences | 1:200 |
| CD45.1 | Flow cytometry | PE | A20 | 1153540 | Sony Biotechnology | 1:200 |
| CD19 | Flow cytometry | PE-Cy7 | eBio1D3 (1D3) | 25-0193-82 | Thermo Fisher Scientific | 1:200 |
| CD11b | Flow cytometry | APC | M1/70 | 1106060 | Sony Biotechnology | 1:800 |
| CD3 | Flow cytometry | Alexa Fluor® 700 | 17A2 | 1101080 | Sony Biotechnology | 1:400 |
| CD45.1 | Lineage depletion | Biotin | A20 | 110703 | Biolegend | 1:300 |
| TER-119 | Lineage depletion | Biotin | TER-119 | 1181020 | Sony Biotechnology | 1:300 |
| CD4 | Lineage depletion | Biotin | GK1.5 | 1102020 | Sony Biotechnology | 1:300 |
| CD8a | Lineage depletion | Biotin | 53-6.7 | 1103520 | Sony Biotechnology | 1:300 |
| B220 | Lineage depletion | Biotin | RA3-6B2 | 1116020 | Sony Biotechnology | 1:300 |
| CD16/32 | Lineage depletion | Biotin | 93 | 101303 | BioLegend | 1:300 |
| CD41 | Lineage depletion | Biotin | MWReg30 | 133930 | BioLegend | 1:300 |
| Sca-1 | Lineage depletion | Biotin | D7 | 108103 | BioLegend | 1:300 |
| Gr-1 | Lineage depletion | Biotin | RB6-8C5 | 1142020 | Sony Biotechnology | 1:300 |
| CD11b | Lineage depletion | Biotin | M1/70 | 1106020 | Sony Biotechnology | 1:300 |
| EIF6 | Immunoblot | | | GTX117971 | GeneTex | 1:1000 |
| GAPDH | Immunoblot | | | G9545 | Merck | 1:10000 |
| eS19 (Rps19) | Immunoblot | | | 15085-1-AP | Proteintech | 1:1000 |
| Anti-rabbit IgG | Immunoblot | HRP | | 7074 | Cell Signaling Technology | 1:5000 |

**Supplementary Table 2. Gene Set Enrichment Analysis of RNA-seq data: A list of positively enriched MSigDB Hallmark gene sets**

| NAME | NES | NOM p-val | FDR q-val |
|---|---|---|---|
| HALLMARK_MYC_TARGETS_V1 | 3.0671105 | 0.0 | 0.0 |
| HALLMARK_E2F_TARGETS | 2.9481921 | 0.0 | 0.0 |
| HALLMARK_OXIDATIVE_PHOSPHORYLATION | 2.8290706 | 0.0 | 0.0 |
| HALLMARK_G2M_CHECKPOINT | 2.642201 | 0.0 | 0.0 |
| HALLMARK_MYC_TARGETS_V2 | 2.5719438 | 0.0 | 0.0 |
| HALLMARK_MTORC1_SIGNALING | 2.0195954 | 0.0 | 2.4509805E-4 |
| HALLMARK_DNA_REPAIR | 1.972006 | 0.0 | 2.1008404E-4 |
| HALLMARK_FATTY_ACID_METABOLISM | 1.9347863 | 0.0 | 1.8382353E-4 |
| HALLMARK_P53_PATHWAY | 1.7502347 | 0.0 | 0.0028431618 |
| HALLMARK_NOTCH_SIGNALING | 1.7231821 | 0.004950495 | 0.0034880443 |
| HALLMARK_UNFOLDED_PROTEIN_RESPONSE | 1.699769 | 0.0 | 0.0042198314 |
| HALLMARK_ADIPOGENESIS | 1.5583233 | 0.0 | 0.01885677 |
| HALLMARK_WNT_BETA_CATENIN_SIGNALING | 1.5022635 | 0.03307888 | 0.02683837 |
| HALLMARK_UV_RESPONSE_UP | 1.4894588 | 0.0037037036 | 0.027358083 |
| HALLMARK_SPERMATOGENESIS | 1.4173876 | 0.026627218 | 0.045275893 |
| HALLMARK_GLYCOLYSIS | 1.3756042 | 0.008333334 | 0.05903587 |
| HALLMARK_XENOBIOTIC_METABOLISM | 1.2991815 | 0.021201413 | 0.096688464 |
| HALLMARK_ESTROGEN_RESPONSE_LATE | 1.2183263 | 0.07272727 | 0.16246332 |
| HALLMARK_APICAL_SURFACE | 1.1347575 | 0.2962963 | 0.27280608 |
| HALLMARK_APOPTOSIS | 1.1281884 | 0.18505338 | 0.27155006 |
| HALLMARK_ESTROGEN_RESPONSE_EARLY | 1.128132 | 0.18181819 | 0.25871834 |
| HALLMARK_REACTIVE_OXYGEN_SPECIES_PATHWAY | 1.1118554 | 0.30027547 | 0.27313113 |
| HALLMARK_PI3K_AKT_MTOR_SIGNALING | 1.0424879 | 0.36482084 | 0.3931973 |
| HALLMARK_MITOTIC_SPINDLE | 0.92531127 | 0.72772276 | 0.66031086 |
| HALLMARK_BILE_ACID_METABOLISM | 0.9169759 | 0.6 | 0.65465826 |

**Supplementary Table 3. Gene Set Enrichment Analysis of RNA-seq data: A list of negatively enriched MSigDB Hallmark gene sets**

| NAME | NES | NOM p-val | FDR q-val |
|---|---|---|---|
| HALLMARK_INFLAMMATORY_RESPONSE | -2.0337608 | 0.0 | 0.0 |
| HALLMARK_COAGULATION | -1.8884282 | 0.0 | 0.0017206814 |
| HALLMARK_COMPLEMENT | -1.5281032 | 0.012396694 | 0.07696216 |
| HALLMARK_TGF_BETA_SIGNALING | -1.51722 | 0.023696683 | 0.06397276 |
| HALLMARK_EPITHELIAL_MESENCHYMAL_TRANSITION | -1.4826974 | 0.011477762 | 0.073612124 |
| HALLMARK_IL2_STAT5_SIGNALING | -1.4714969 | 0.00807537 | 0.06855643 |
| HALLMARK_KRAS_SIGNALING_UP | -1.4223773 | 0.023319615 | 0.09558294 |
| HALLMARK_UV_RESPONSE_DN | -1.402376 | 0.03806735 | 0.100769944 |
| HALLMARK_HYPOXIA | -1.3578143 | 0.045081966 | 0.13083611 |
| HALLMARK_IL6_JAK_STAT3_SIGNALING | -1.3221637 | 0.1017192 | 0.1573019 |
| HALLMARK_TNFA_SIGNALING_VIA_NFKB | -1.3047135 | 0.048209365 | 0.16406558 |
| HALLMARK_KRAS_SIGNALING_DN | -1.3038148 | 0.09090909 | 0.15133482 |
| HALLMARK_ALLOGRAFT_REJECTION | -1.2548201 | 0.10339943 | 0.2062244 |
| HALLMARK_ANGIOGENESIS | -1.2407737 | 0.2 | 0.21205784 |
| HALLMARK_INTERFERON_GAMMA_RESPONSE | -1.1463896 | 0.20526315 | 0.3597538 |
| HALLMARK_CHOLESTEROL_HOMEOSTASIS | -1.1295773 | 0.281106 | 0.37050083 |
| HALLMARK_MYOGENESIS | -1.025393 | 0.40143886 | 0.5846858 |
| HALLMARK_INTERFERON_ALPHA_RESPONSE | -1.0191847 | 0.43472022 | 0.56760556 |
| HALLMARK_PROTEIN_SECRETION | -0.9452486 | 0.54572713 | 0.71615493 |
| HALLMARK_APICAL_JUNCTION | -0.81811005 | 0.82865167 | 0.9811232 |
| HALLMARK_HEDGEHOG_SIGNALING | -0.80280477 | 0.73504275 | 0.9628326 |
| HALLMARK_HEME_METABOLISM | -0.79222816 | 0.89747006 | 0.9369517 |
| HALLMARK_ANDROGEN_RESPONSE | -0.68855524 | 0.96253604 | 1.0 |
| HALLMARK_PEROXISOME | -0.64410555 | 0.9769784 | 0.98277134 |

**Supplementary Table 4. A list of RT-qPCR primers used in this study**

| Gene | Primer |
|---|---|
| *Actb* | CCACAGCTGAGAGGCAAATC |
| | CTTCTCCAGGGAGGAAGAGG |
| *Eif6* | CCAAGTACCATTGCCACCAG |
| (mouse) | GGAAAATGAGCCAAAGTCCAGAG |
| *Eif6 + EIF6* | AATGTCACCACCTGCAATGAC |
| (mouse + transgene) | TGTCTGAAGACTTCCACCTTGAG |
| *Rps19* | GCAGAGGCTCTAAGAGTGTGG |
| | CCAGGTCTCTCTGTCCCTGA |
| *Cybb* | CTTTCTCAGGGGTTCCAGTG |
| | TGCAGTGCTATCATCCAAGC |
| *S100a9* | CTGACACCCTGAGCAAGAAG |
| | CCTGGTTTGTGTCCAGGTC |

### SUPPLEMENTARY FIGURE 1

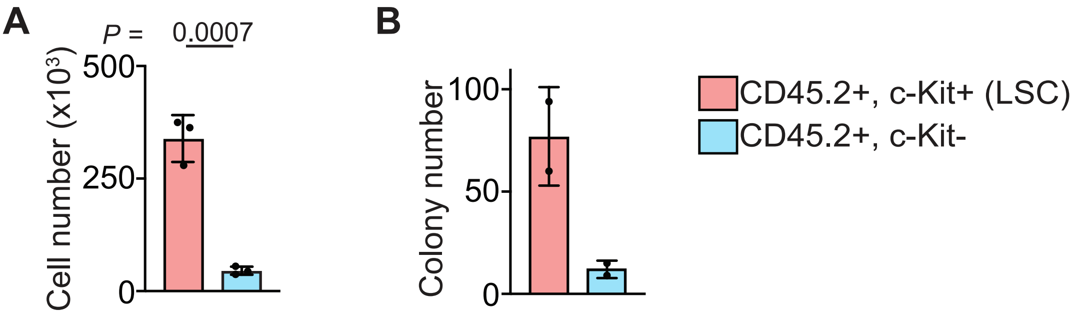

### SUPPLEMENTARY FIGURE 2

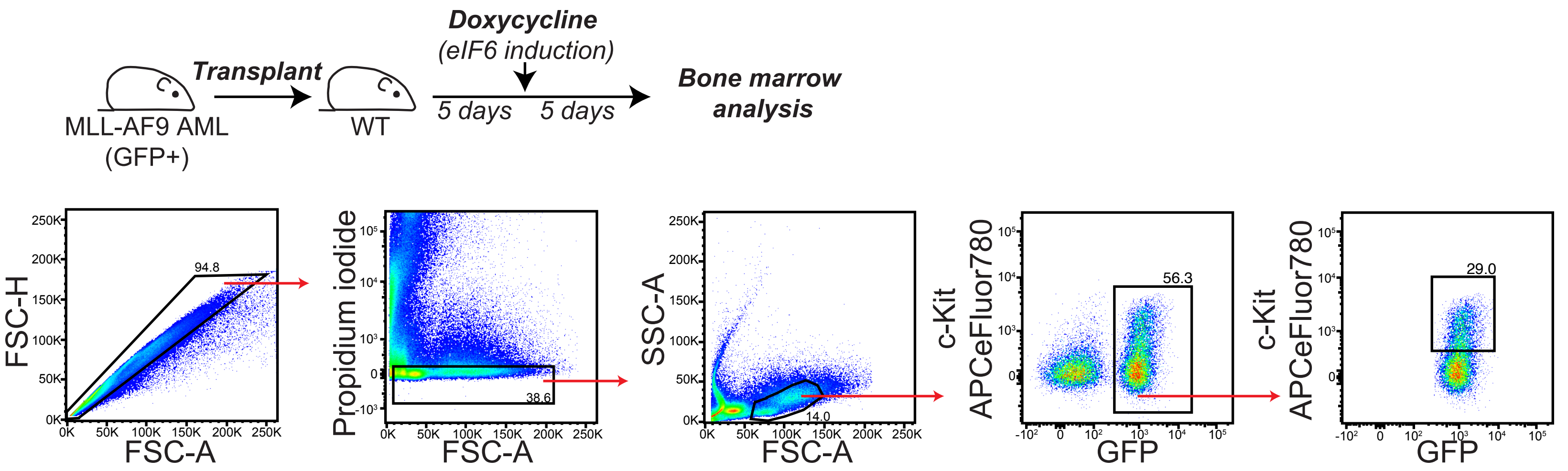

### SUPPLEMENTARY FIGURE 3

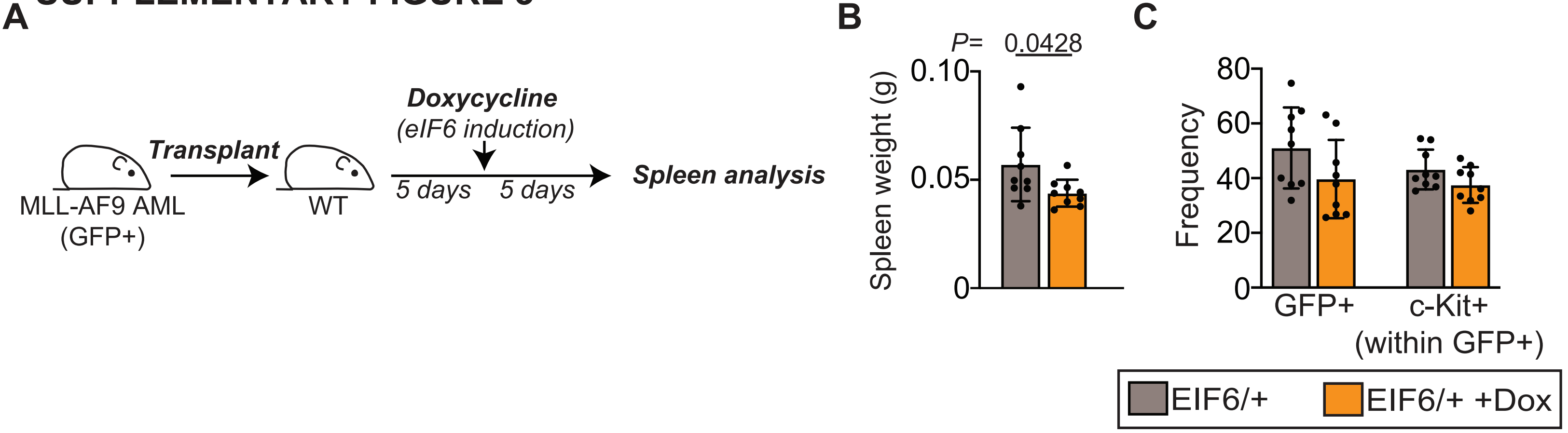

SUPPLEMENTARY FIGURE 4

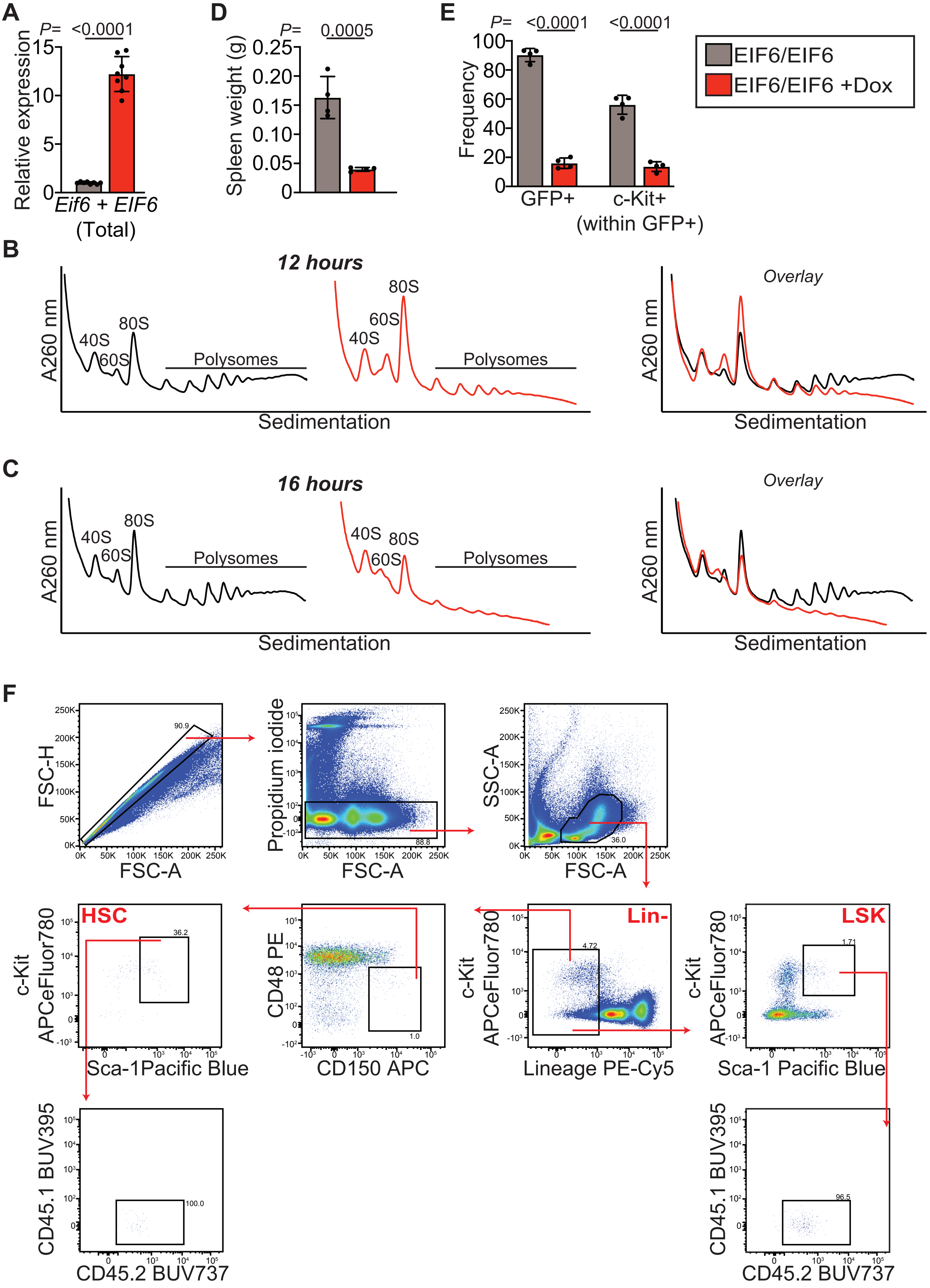

#### SUPPLEMENTARY FIGURE 5

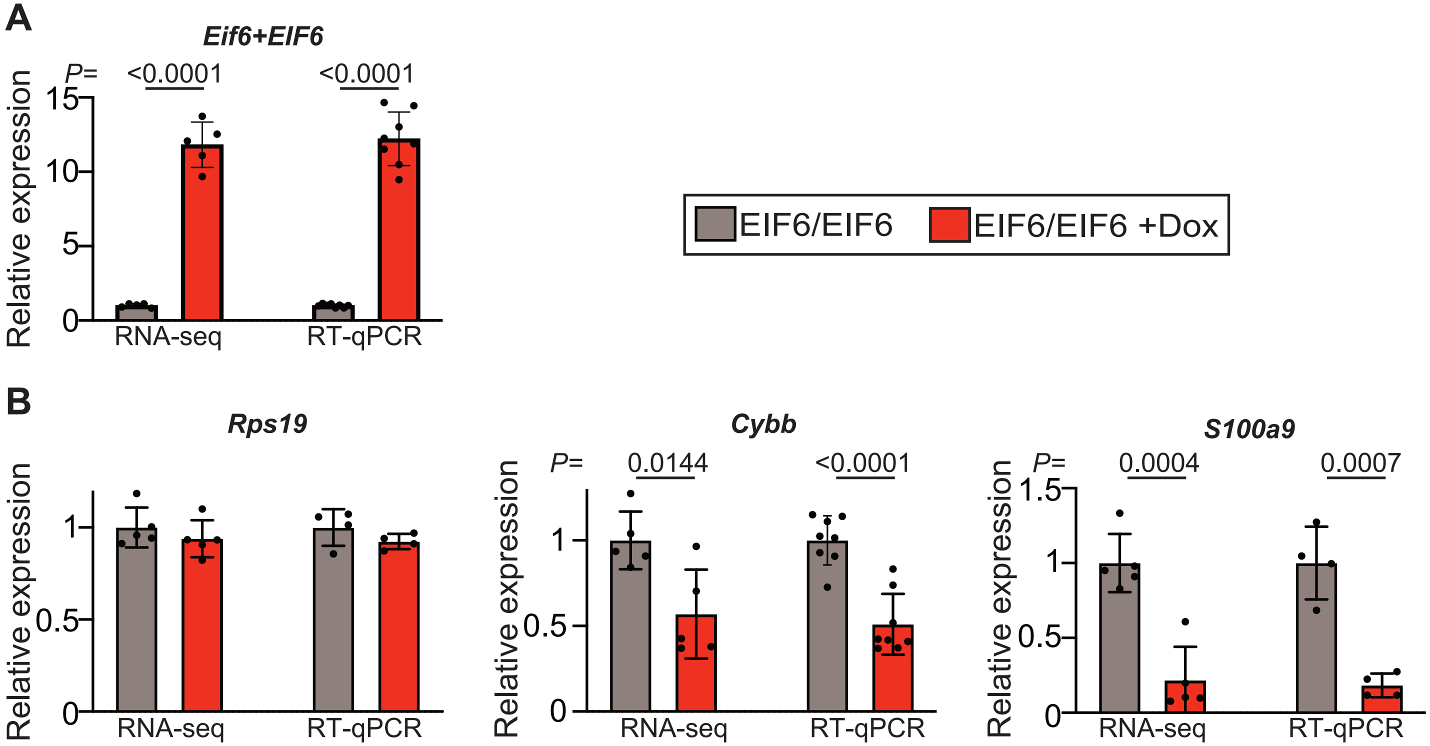

### SUPPLEMENTARY FIGURE 6

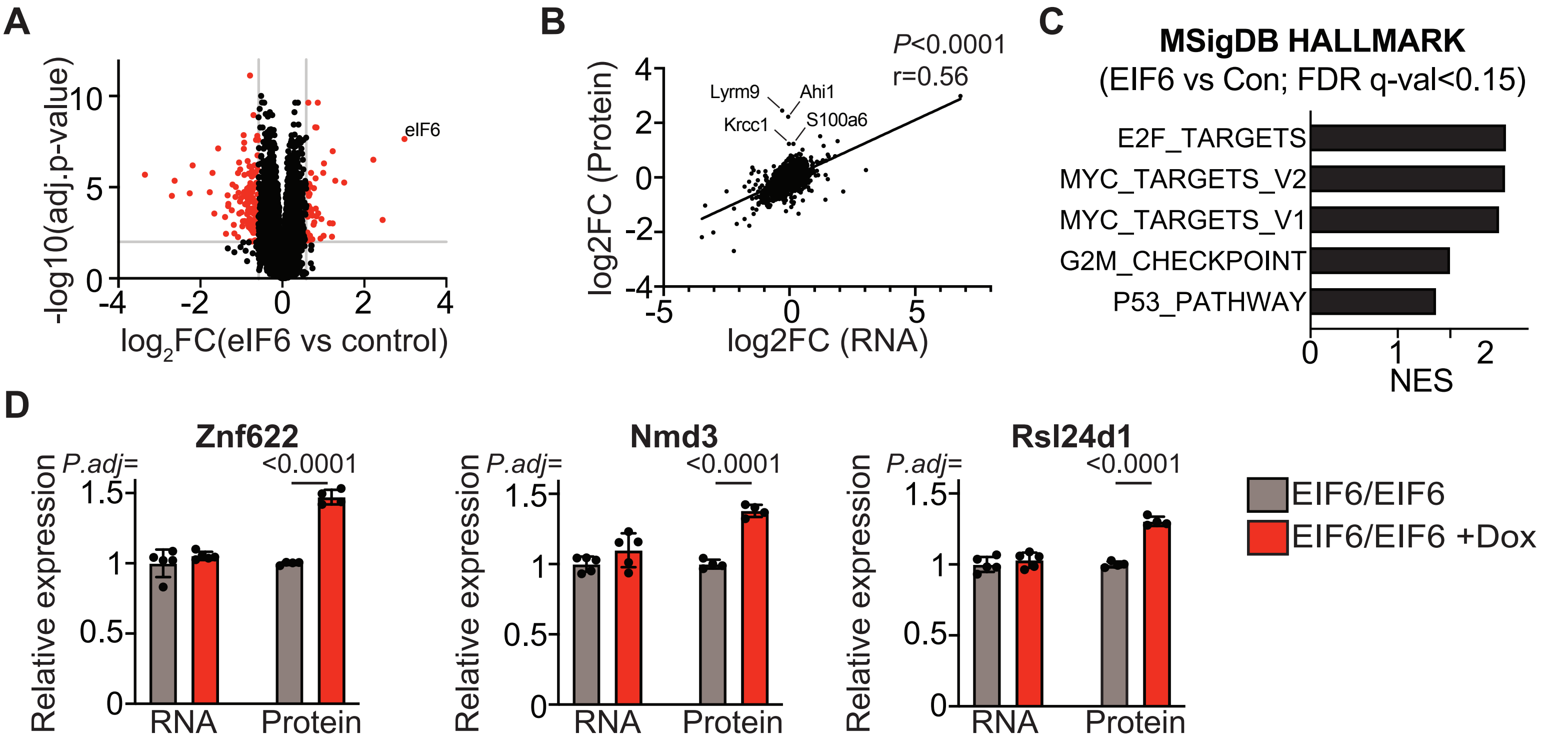

### SUPPLEMENTARY FIGURE 7

**A**

```
Reference:    5'-TTATTCTTGCTCTTAGGCCTGGCTCCTCCCAGCATCTTATCCGGGTGGAAGGAAATTTG-3'
Allele 1      5'-TTATTCTTGCTCTTAGGCCTGGCTCCTC-------TCTTATCCGGGTGGAAGGAAATTTG-3'
Allele 2      5'-TT--------------------------------------CCGGGTGGAAGGAAATTTG-3'
```

Exon 6
PAM
gRNA

**B**

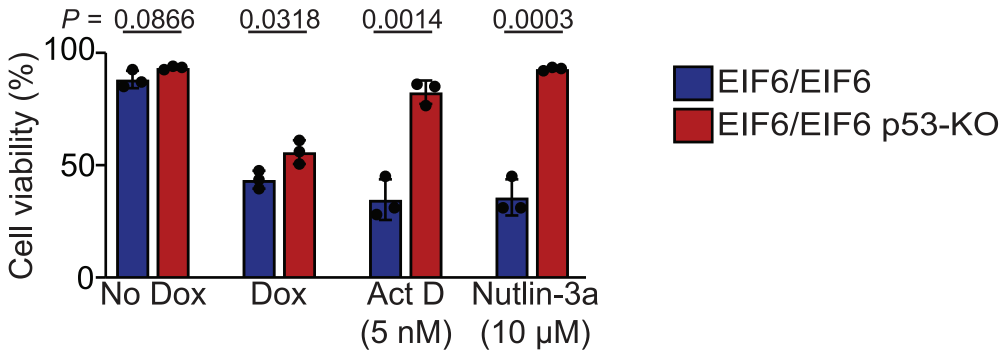
